# Rapid fluorescence lifetime sensor development of LifeCamp enables transient and baseline absolute calcium measurements

**DOI:** 10.64898/2025.12.23.696288

**Authors:** Bart Lodder, Michelle Raghubardayal, Sanika Ganesh, Xintong Cai, Josh Stern, Michael Sherman, Paul Rosen, Tarun Kamath, Isa Hartman, Michelle Siegel, Joshua Timmins, Roger Adan, Mark Andermann, Bernardo L. Sabatini

## Abstract

Genetically encoded calcium sensors (GECIs) have been instrumental for studying neuronal activity and intracellular signaling. GECIs are typically fluorescence-intensity sensors that change brightness upon calcium binding. Iterative improvements in GECIs have yielded indicators that report action potential-evoked calcium entry with high sensitivity and temporal resolution, enabling measurement of network activity across thousands of neurons. However, fluorescence intensity-based measurements generally cannot report baseline or absolute calcium levels and may confound neuromodulatory regulation of calcium handling with changes in action potential firing. Fluorescence lifetime sensors are insensitive to many artifacts that plague intensity-based measures and report absolute substrate levels, including those at rest. However, relatively few lifetime sensors for neuronal signals exist, and developing new sensors is typically difficult and low-yield. Here, we introduce a new rapid lifetime sensor development (RALISED) platform, which we use to build a new GCaMP8m-based high-speed lifetime calcium sensor, termed LifeCamp. We show that LifeCamp enables comparison of baseline calcium signals in cell culture, brain slices, and mice. In addition, we show that LifeCamp enables the detection of fast action potential-evoked calcium transients in single neurons from brain slices and in behaving mice. Using LifeCamp, we discovered calcium baseline changes associated with neuronal activity in brain slices and behaving mice, as well as slow average calcium changes in neuronal populations of freely moving mice. Altogether, this study highlights the utility of the RALISED method to rapidly develop new lifetime sensors and the application of the LifeCamp calcium lifetime sensor to study fast and slow calcium signaling.

**Significance statement:** We developed a new high-speed, sensitive calcium lifetime sensor (LifeCamp) using a novel rapid lifetime sensor development (RALISED) platform. LifeCamp has high sensitivity and a large substrate-dependent lifetime change (<1ns), allowing for the capture of baseline calcium levels, transient calcium changes, and neuronal firing in vitro and behaving animals. LifeCamp lifetime measurement is insensitive to artifacts that plague conventional intensity imaging and enables absolute comparison of baseline and transient calcium changes across cells, brain regions, and experimental conditions. Hence, LifeCamp is a powerful tool that enables a more accurate and in-depth understanding of neuronal activity and calcium signaling.

## Introduction

The development of genetically encoded fluorescent indicators (GEFIs) for a wide range of molecules has been transformative for the study of neuronal signaling. GEFIs can be expressed in specific cell types and enable the detection of neuronal signals, such as neurotransmitters, transmembrane voltages, and intracellular signals, in brain slices and behaving animals (1–4). Upon substrate binding, GEFIs typically change their brightness, which can be detected and used to measure relative substrate changes (5–7). A frequently used subcategory of GEFIs are the genetically encoded calcium indicators (GECIs) (2). Calcium (Ca) is a ubiquitous second messenger whose concentration changes in response to neuronal firing, G-protein coupled receptor activation, synaptic activity, and metabolic changes, and, in turn, regulates plasticity, neuronal excitability, neurotransmitter release, gene expression, and mitochondrial function (5). In neuroscience, high-speed GECIs, which have taken decades to develop, are used to detect action potential firing of many individual neurons in brain slices and model organisms using fluorescence microscopy (2, 6). In addition, measurement of GECIs using fiber photometry reports changes in bulk Ca levels in selected neuronal populations in freely moving animals, which are used as a proxy for neuronal activity dynamics (7–9).

Although currently available GECIs have contributed greatly to our understanding of the brain, several limitations remain. Currently used high-speed GECIs report Ca levels by changing their brightness, which is read out as changes in fluorescence intensity. However, fluorescence intensity measurements are sensitive to sensor expression and hemodynamic, motion, and photobleaching artifacts, which can affect the fluorescence intensity signal on short (ms-s) and long (min-hr) timescales (6, 9–14). These artifacts are difficult to correct for and distinguish from biological signals (6, 9–14). Therefore, fluorescence intensity measurements typically limit GECI application to the detection of relative and transient Ca levels and preclude the detection of resting Ca levels and comparisons of absolute Ca levels between cell types and brain regions (10, 15–17).

Ideally, artifact-insensitive and high-speed measurements of a biological signal could be made in absolute units over long timescales (min-days). Fluorescence lifetime (i.e., the time it takes for an excited fluorophore to release a photon after excitation) can be used to detect substrate changes as an alternative to fluorescence intensity. Fluorescence lifetime is an intrinsic property of each fluorophore molecule and thus independent of GEFI concentration (as long as autofluorescence is negligible compared to sensor fluorescence) and insensitive to many of the artifacts that limit intensity measurements, such as hemodynamic-, motion-, and photobleaching-related artifacts(11, 14, 15, 18) (although see (19)). Some GEFIs, e.g., the dopamine sensor dLight3.8 and acetylcholine sensor GRAB-Ach, change fluorescence lifetime upon substrate binding (15, 20, 21). These sensors, together with the recent development of high-speed fluorescence lifetime microscopy (FLIM) and fluorescence lifetime photometry (FLIP) systems, now allow for artifact-insensitive absolute measurement of transient and baseline changes in neuronal signals(15, 18, 22–24).

However, the application of lifetime systems and the investigation of fast and slow absolute neuronal signals are limited by the relatively few fluorescence lifetime sensors available (15). For example, no high-speed Ca lifetime sensor exists that allows for both measurement of baseline Ca levels and action potential rates in mice. Molecular mechanisms that cause lifetime change in GECIs are not yet fully understood, and it is still difficult to model and develop new lifetime sensors in a rational manner(25, 26). Therefore, the development of lifetime sensors typically requires high-throughput screening of many constructs, which is difficult, costly, and time-consuming(27). This limits the development of new lifetime sensors and subsequent biological discoveries.

Here, we describe a new rapid lifetime sensor development (RALISED) platform that allows for rational, fast, and affordable lifetime sensor development. The core concept is that existing intensity sensors can be transformed into lifetime sensors by fusion with a fluorescent protein variant of an appropriate lifetime. By linking the intensity sensor GCaMP8m(6) to ultra-short lifetime green fluorescent protein BrUSLEE(28), we developed a new green high-speed Ca lifetime sensor, termed LifeCamp. We show that LifeCamp has a large (>1ns) substrate-dependent lifetime change and enables fast measurement of baseline and spiking-related Ca signals *in vitro* and *in vivo*. We show that lifetime LifeCamp measurements are insensitive to intensity artifacts and allow accurate interpretation of measured signals. We applied LifeCamp for biological discovery and studied the relationship between neuronal firing and baseline Ca, characterized baseline Ca levels across brain regions, and discovered novel slow baseline Ca changes in neurons of behaving mice.

## Results

### Rapid lifetime sensor development (RALISED) of LifeCamp

To develop a high-speed Ca lifetime sensor in a cost-efficient, rapid manner, we created a new rapid lifetime sensor development platform, termed RALISED. RALISED enables rapid lifetime sensor development by building on existing intensity sensors. Intensity sensors are optimized to have a large intensity change between the substrate-bound and unbound states, but are not designed or expected to have a lifetime change. Using RALISED, lifetime sensors are created by linking existing intensity sensors to static fluorophores with the same spectral properties but a different lifetime. Measurements of fluorescence lifetime from the fusion protein reflect contributions from both fluorophores and equal the intensity-weighted average of the lifetimes of each. The measured average lifetime (*τObserved*) is determined by the sum of the intensity contribution of the intensity sensor (*I1*) and the static fluorophore (*I2*), multiplied by their respective lifetimes (*τ1*, *τ2*), and together divided by the total observed intensity (*Itotal*) (Fig. 1*A*, Eq. 1):

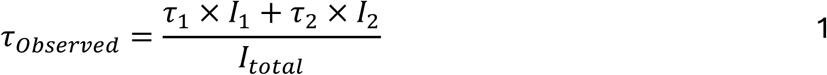

**Fig. 1:**
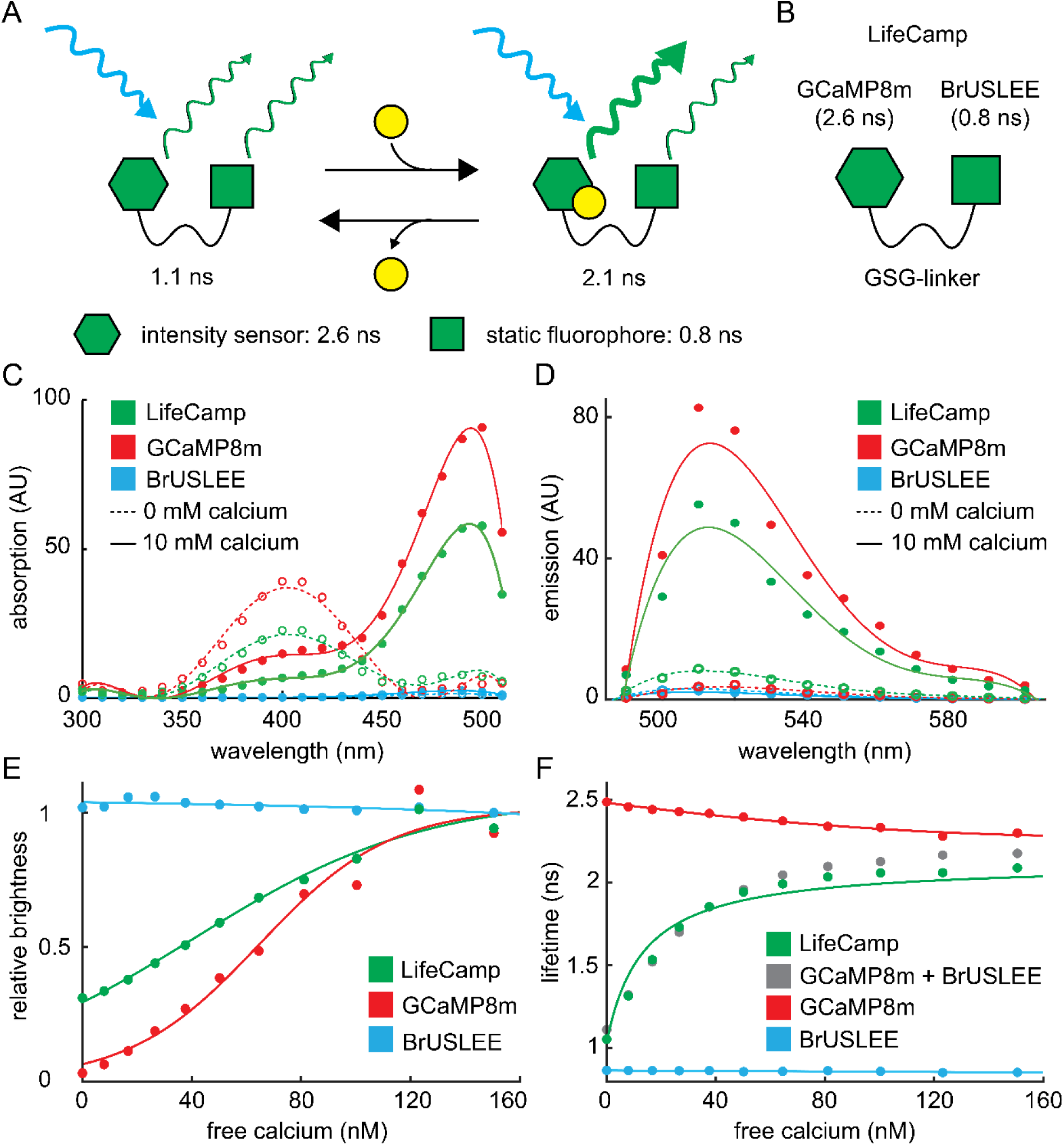
RALISED designed sensor LifeCamp and purified protein characterization. a, Schematic overview of RALISED sensors with example construct in unbound (left) and substrate-bound state (right). **b,** Schematic overview of LifeCamp and the selected fluorophores, GCaMP8m and BrUSLEE, with their fluorescent lifetimes, and linked using a GSG linker. **c,** Absorption spectra of LifeCamp, GCaMP8m and BrUSLEE purified protein exposed to 0mM [Ca], 30mM MOPS (dotted line) or 5mM [Ca], 30mM MOPS (solid line) at pH 7.2 at room temperature. Absorption spectra were measured with a center wavelength of 540nm emission light. Line=polynomial fit **d,** Emission spectra of LifeCamp, GCaMP8m and BrUSLEE purified protein exposed to 0mM [Ca], 30mM MOPS (dotted line) or 4mM free [Ca], 30mM MOPS (solid line) at pH 7.2 at room temperature. Emission spectra were measured with a center wavelength of 460nm excitation light. Line=polynomial fit **e,** Relative brightness of LifeCamp, GCaMP8m and BrUSLEE. Constructs were exposed to different free calcium concentrations at pH 7.2 at room temperature. Line=polynomial fit **f,** Fluorescence lifetime measurement of LifeCamp, GCaMP8m and BrUSLEE (mixed and without linker), GCaMP8m and BrUSLEE. Constructs were exposed to different free calcium concentrations at pH 7.2 at room temperature. Lifetime was measured with 480nm (Δλ=10 nm) excitation and 500nm (Δλ=20 nm) excitation light. Red and blue line: polynomial fit, green line: model fit (methods), Kd=126 nM, ΔF=83.25, τGCaMP8m_bound_=2.25 ns, τGCaMP8m_unbound_=2.45 ns, τBrUSLEE=0.86 ns.

Thus, in the unbound state, the intensity sensor is typically dim, resulting in a relatively large contribution of photons from the static fluorophore, leading to an average lifetime closer to the lifetime of the static fluorophore. Conversely, in the bound state, the intensity sensor is bright, resulting in a relatively large contribution of photons from the intensity sensor, leading to an average lifetime closer to the lifetime of the intensity sensor. A sample typically contains many sensors; therefore, in the case of RALISED sensors, the measured lifetime is determined by the ratio of bound and unbound state fluorophores, their respective lifetimes, and intensity contributions (Eq. 1). Changes in substrate concentration and receptor binding lead to a change in the ratio between bound and unbound sensor concentration and change the measured average lifetime. The measurement is inherently ratiometric, rendering it insensitive to changes in RALISED sensor concentration, fluorescence excitation and collection efficiencies, and many other substrate-independent factors.

For optimal RALISED sensors, the intensity sensor and static fluorophore components should have overlapping excitation and emission spectra, comparable bleaching rates, and similar brightness. Optimal RALISE sensors can be developed by selecting intensity sensors with a large intensity change upon substrate binding, and by selecting the static fluorophore based on maximizing the difference in the static and intensity fluorophore lifetimes. For ligand-sensitive RALISED sensors, the apparent affinity (i.e., the midpoint of the dose-response curve is determined by the affinity of the intensity sensor and the substrate-dependent intensity change of the RALISED sensor (methods). Therefore, the sensitivity of the fluorescence measurement can differ substantially between lifetime and intensity modes.

To develop a Ca lifetime sensor, we selected the intensity Ca sensor GCaMP8m, given its high-speed properties, stability in biological systems, and its large intensity change between bound and unbound states (6). We identified the GFP variant “bright ultimately short lifetime enhanced emitter (BrUSLEE)” as one having a large difference in lifetime (∼0.8 ns) from reported lifetimes of GCaMP8m (∼2.6 ns) and good reported stability *in vivo* (26, 28). We added a flexible GSG linker sequence between GCaMP8m and BrUSLEE to ensure stochiometric expression and colocalization in the sample (Fig. 1*B*).

### Purified protein validation

To validate and characterize LifeCamp *in vitro* as an intensity and lifetime sensor, we determined the excitation and emission spectra as well as intensity and lifetime of LifeCamp, GCaMP8m, and BrUSLEE as a function of [Ca]. We measured the excitation and emission spectra of purified proteins in an intensity plate reader in 0 and 150 nM free [Ca], which confirmed that the excitation and emission spectra of GCaMP8m and LifeCamp depend on [Ca] and that the spectrum of LifeCamp is intermediate to those of BrUSLEE and GCaMP8m (Fig. 1*C* and *D*). Similarly, we observed a large change in intensity for GCaMP8m and LifeCamp, and minimal change for BrUSLEE, across a physiologically relevant range of [Ca] (Fig. 1*E*). We observed minimal changes in lifetime for BrUSLEE and GCaMP8m across [Ca]. In contrast, as expected and by design, we observed a large lifetime change of 1.03 ns for LifeCamp between 0 and 150 nM free [Ca] (Fig. 1*F*, *SI Appendix,* Fig. S1). Furthermore, the lifetime and intensity across [Ca] were nearly identical for LifeCamp and 1:1 mixtures of unlinked GCaMP8m and BrUSLEE purified protein, indicating that the fluorescent properties of the component fluorophores were unperturbed by their fusion via the linker (Fig. 1*F*).

### HEK293T cell validation

To determine the viability of LifeCamp expression and the utility of its dynamic range in eukaryotic cells, HEK293T cells were transfected with LifeCamp, BrUSLEE, and GCaMP8m and imaged using two-photon fluorescence lifetime microscopy (2P-FLIM) (Fig. 2*A*). HEK293T cells were incubated and imaged in buffer solution containing a Ca ionophore and either no free [Ca], to deplete intracellular [Ca], or with a high concentration of [Ca] (10 mM) to increase intracellular [Ca]. We observed that cells expressing LifeCamp had a significant change in lifetime between the 0 mM [Ca] (1.62 ns) and 10 mM [Ca] (2.16 ns; paired t-test, p=2.1E-15, n=12 cells) solutions (Fig. 2*B* and *C*). We also observed a small but significant change in the lifetime between depleted and high intracellular Ca HEK293T cells that were transfected with GCaMP8m (paired t-test, p=4.01E-7, n=12 cells) (Fig. 2*C*). Moreover, we observed that lifetime variance across HEK293T cells at baseline (2 mM [Ca] without Ca^2+^ ionophore) was larger than in conditions in which intracellular [Ca] was clamped by inclusion of a Ca^2+^ ionophore (F-test, p=7E-4, n=12 sites; *SI Appendix,* Fig. S2), indicating that lifetime variance between HEK293T cells reflects differences in intracellular [Ca] across cells.

**Fig. 2:**
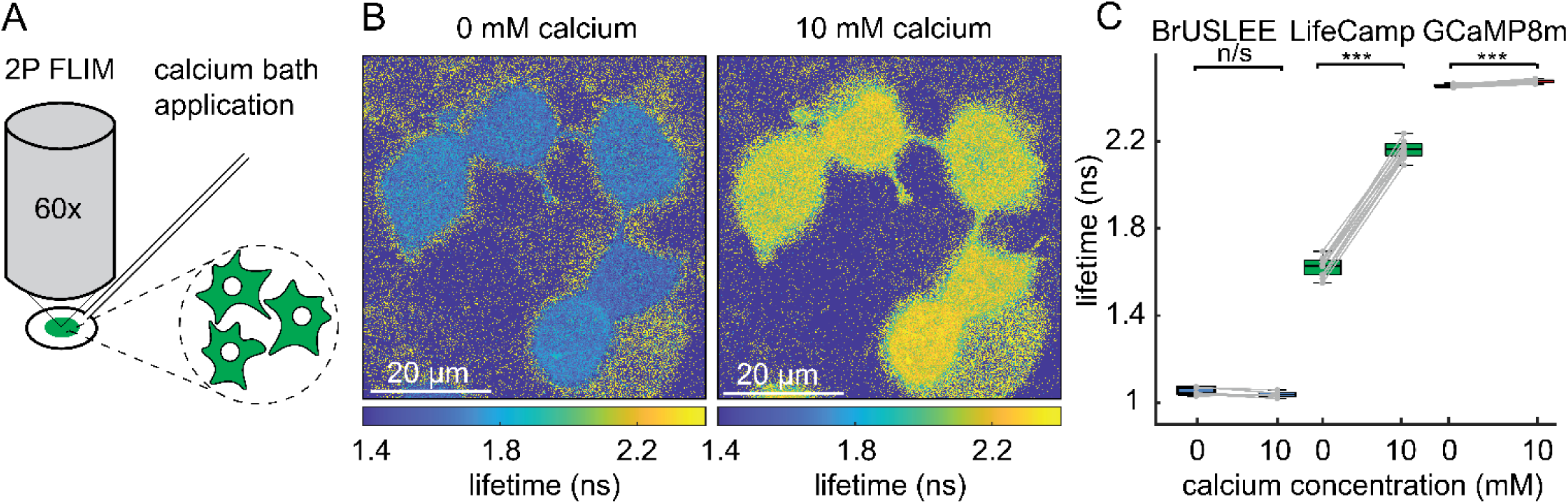
Characterization of LifeCamp in HEK293T cells. a, Schematic of 2P-FLIM setup for imaging LifeCamp, BrUSLEE or GCaMP8m in HEK293T cells. HEK293T cells, transfected with one of the three constructs, were first incubated and imaged in external bath solution with calcium chelating agent ionomycin (5 µM) without [Ca] (depleted condition), and afterwards incubated and imaged in external bath solution with [Ca] (10mM, high calcium condition). **b,** 2P-FLIM lifetime example images show uniform expression of LifeCamp in HEK293T cells in depleted condition (left) and high calcium condition (right). **c,** LifeCamp lifetime was significantly increased (paired t-test, p=2.1E-15, n=12 cells), GCaMP8m lifetime was slightly increased (paired t-test, p=4.01E-7, n=12 cells) and BrUSLEE lifetime did not change (paired t-test, p=0.10, n=5 cells) between the 0 mM and 10 mM calcium conditions. Boxplot elements: center mark, median; box limits, upper and lower quartiles; whiskers, furthest datapoints from the median.

### Neuronal firing measured in brain slices

To investigate the ability of LifeCamp to report the occurrence of single and multiple action potentials in neurons, we expressed LifeCamp or GCaMP8m in the M1 cortex and evoked action potentials using a stimulating electrode placed in the periphery of a neuron in brain slices while simultaneously imaging them using 2P-FLIM (Fig. 3*A*-*C*). Stimulation of LifeCamp-expressing neurons evoked a transient increase in lifetime (Fig. 3*D*-*F*). We stimulated neurons at frequencies of 1 Hz, 5 Hz, 10 Hz, and 20 Hz for 600ms, and we observed a progressively increased lifetime response of LifeCamp. An individual action potential evoked a ∼40 ps change, while firing rate at 20 Hz increased it by > 100 ps. Conversely, we observed minimal lifetime responses for GCaMP8m (Fig. 3*F*). In addition, the intensity characteristics of LifeCamp were intact, even though, as expected, the dF/F was slightly lower for LifeCamp compared to GCaMP8m (Fig. 3*F*). These results show that LifeCamp can be used in both lifetime and intensity modes, depending on the experimental requirements. The high speed of LifeCamp facilitated the identification of individual action potentials fired at high frequency (>10Hz) using both lifetime and intensity mode. We did not observe a change in the kinetics of single-action-potential intensity responses of GCaMP8m and LifeCamp, again indicating that GCaMP8m properties in LifeCamp are unaffected.

**Fig. 3:**
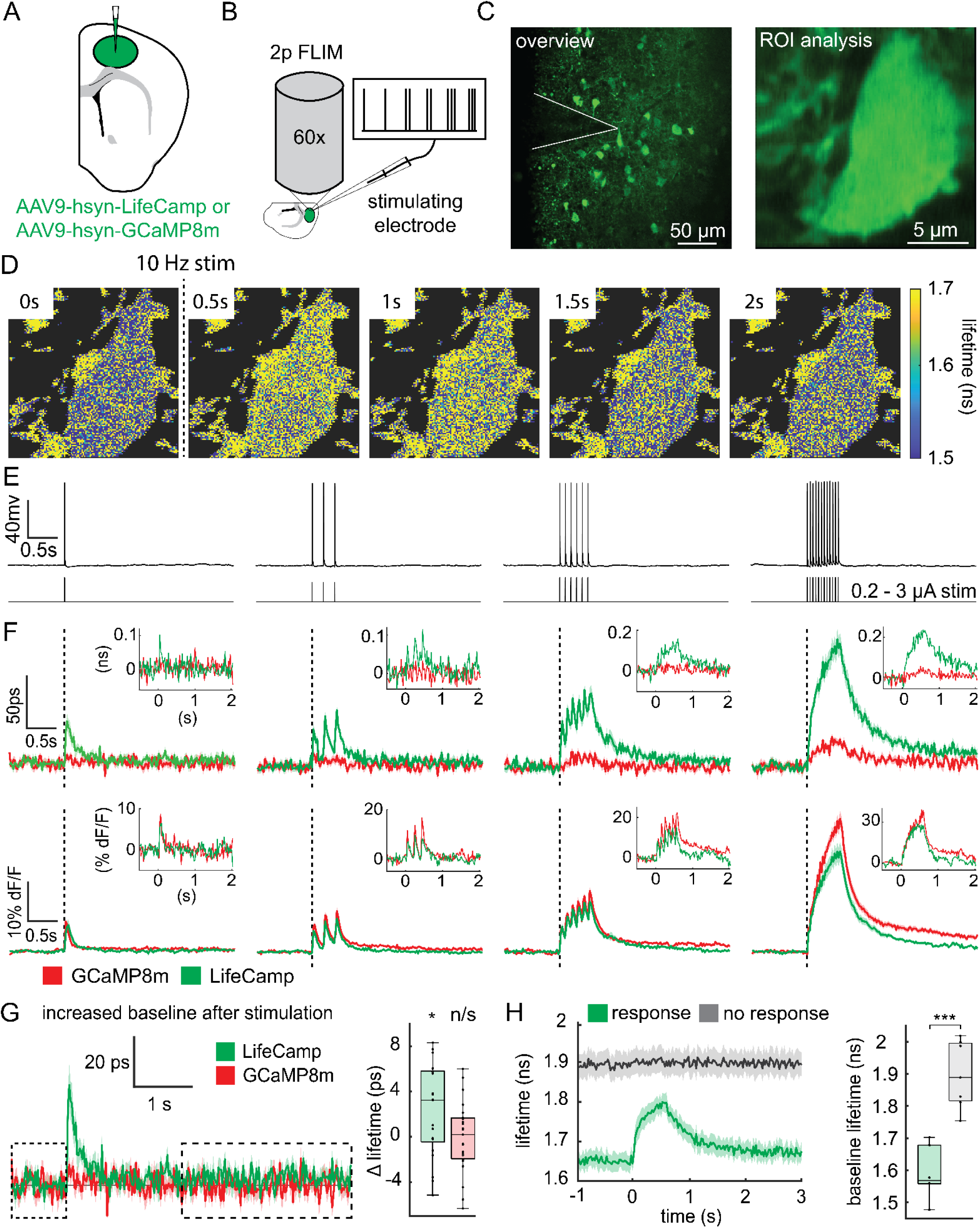
LifeCamp measurements of stimulated neuronal firing. a, Schematic of coronal section and viral injection of AAV9-hsyn-LifeCamp or AAV9-hsyn-GCaMP8m into the M1 cortex of a mouse. **b,** Schematic of 2P-FLIM setup for imaging and stimulation of LifeCamp expressing M1 cortical neurons. **c,** Overview image with stimulating electrode on top of LifeCamp expressing M1 cortical neuron (left) and example of ROI used for analysis (right). **d,** Frames captured of an example neuron stimulated at 10 Hz with the stimulating electrode. **e,** Whole cell recordings of a neuron stimulated using firing patterns (1, 5, 10, 20 Hz, left to right) as used during 2P-FLIM recordings. **f,** Fluorescence lifetime (top) and intensity (bottom) recordings of LifeCamp averaged across trials: (1 Hz n=21, 5 Hz n=24, 10 Hz n=22 and 20 Hz n=26 trials from 7 cells, 3 mice). GCaMP8m: (1 Hz n=23, 5 Hz n=24, 10 Hz=25 and 20 Hz n=23 trials from 6 cells, 3 mice). Example trials are displayed in the top right corner of each graph. Error bar=SEM. **g,** Increased baseline fluorescence lifetime after 1 Hz stimulation (p=0.0249, paired t-test, n=7 cells, 3 mice). No observed change in fluorescence lifetime was observed for GCaMP8m (p=0.9376, paired t-test, n=6 cells, 3 mice). **h,** Baseline lifetime measured in non-responding and responding cells (p=9.21E-5, two sample t-test, n=8 cells, 4 animals not responding, n=7 cells, 3 animals). Trace is aligned to stimulation at 0s. Boxplot elements: center mark, median; box limits, upper and lower quartiles; whiskers, furthest datapoints from the median.

Using LifeCamp, we observed a significant increase in baseline Ca lasting several seconds after single action potential firing (p=0.0249, paired t-test, n=7 cells, 3 mice) (Fig. 3*G*). We did not observe a significant change in the lifetime of GCaMP8m in response to action potential firing (p=0.9376, paired t-test, n=6 cells, 3 mice).

Not all neurons responded to electrical stimulation. Interestingly, we observed that LifeCamp expressed in responsive neurons had a mean lifetime of 1.55-1.72 ns. In contrast, unresponsive neurons had a lifetime of 1.77-2.01 ns (p=9.21E-5, two-sample t-test, n=8 cells, 4 animals not responding, n=7 cells/3 mice) (Fig. 3*H*). In addition, we observed several instances in which, during stimulation, neurons dramatically increased Ca levels, whereafter they became unresponsive (*SI Appendix*, Fig. S3). Thus, it is likely that neurons with a high LifeCamp lifetime have high intracellular [Ca], consistent with being unhealthy or damaged during brain slicing. These results indicate that LifeCamp basal lifetime can be used to screen for healthy and unhealthy cells during electrophysiological experiments, something which is not possible based on the brightness of neurons expressing intensity GECIs.

### LifeCamp measurements in freely behaving mice

To determine whether LifeCamp can be utilized to measure [Ca] in neuronal populations *in vivo*, LifeCamp was expressed in dopaminergic neurons (DANs) of the ventral tegmental area (VTA). LifeCamp lifetime was measured in freely moving mice through a fiber optic using fluorescence lifetime photometry at high temporal resolution (FLIPR) (Fig. 4*A*-*C*, *SI Appendix*, Fig. S4)(10). DANs of the VTA fire tonically (0.5-10 Hz) and respond to rewarding stimuli with burst firing (>15 Hz)(30, 31). Thus, to investigate whether LifeCamp can detect fast [Ca] changes *in vivo*, we gave freely moving mice chocolate pellets while simultaneously recording LifeCamp using FLIPR (Fig. 4*C*). We measured large changes in intensity (dF/F=14%) and lifetime (Δτ=70 ps) in response to chocolate pellet consumption (Fig. 4*D*). To control for artifacts, we developed a Ca-insensitive binding mutant for LifeCamp, termed LifeCamp0. LifeCamp0 was developed by replacing the calmodulin binding site in LifeCamp with a 5x GSG linker, which, as previously shown, diminishes Ca-dependent changes in fluorescence (6, 32). The LifeCamp0 intensity increased (dF/F=4%) during chocolate pellet consumption, whereas the lifetime was unaffected, consistent with the relative insensitivity of fluorescence lifetime to artifacts.

**Fig. 4:**
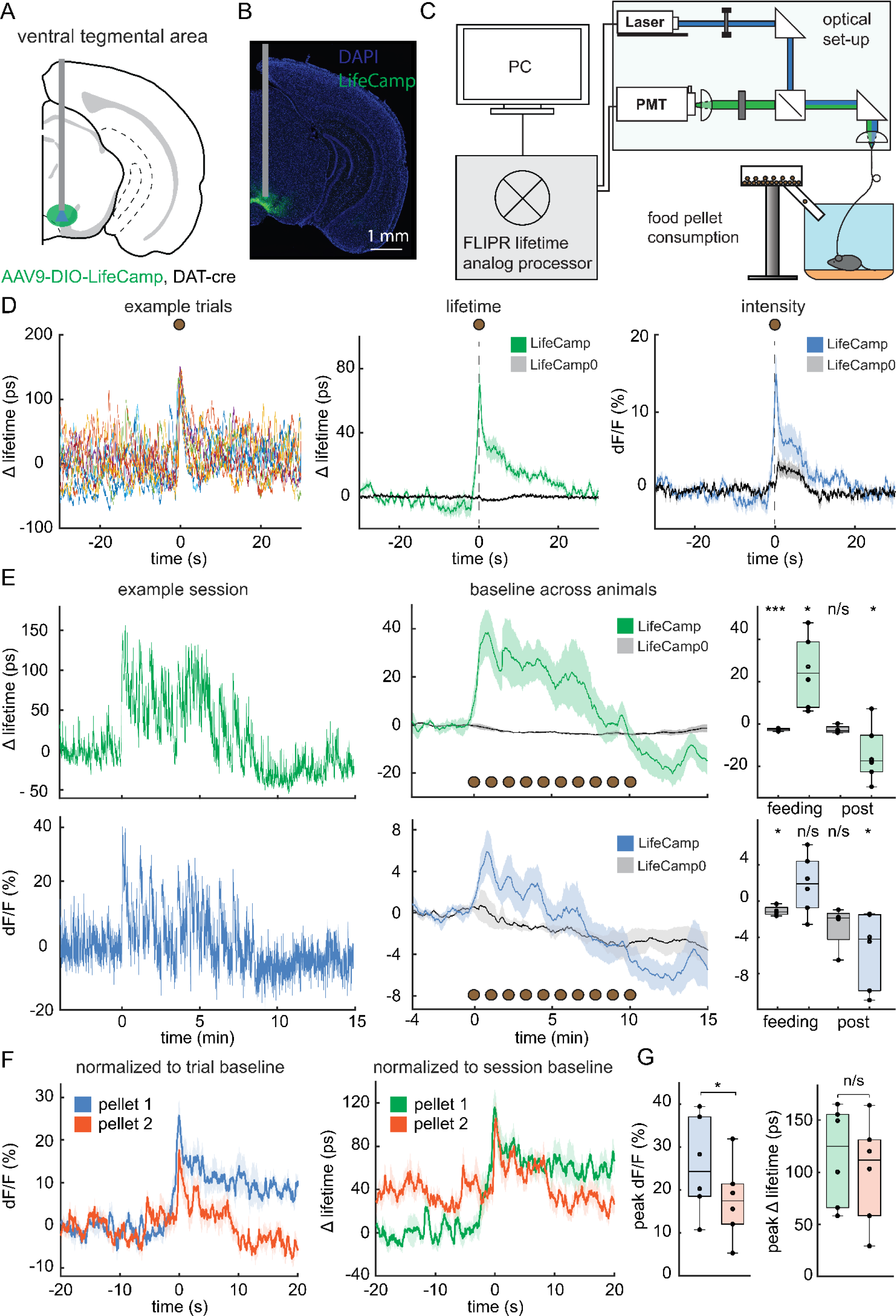
LifeCamp measured in the VTA of freely moving mice with FLIPR. a, Schematic of LifeCamp expression in the VTA of DAT-cre mice. A fiber optic was placed above the injection site, and the fluorescence lifetime was measured using FLIPR in freely moving mice. **b,** Example immunohistochemistry staining of LifeCamp and nuclei (DAPI) to confirm the injection location, expression and fiber optic location in the VTA. **c,** Schematic of the FLIPR lifetime analog processor (left) and optical set-up (right) in combination with behavioral tasks (bottom). Mice received chocolate pellets while freely moving. **d,** VTA DAN LifeCamp lifetime and intensity response to chocolate pellet consumption. Lifetime measurements during example trials of a single animal show an increase in delta lifetime during reward consumption (left). Averaged LifeCamp measurements (n=6 mice) during food pellet consumption show a large change in delta lifetime (middle) and intensity (right). Control sensor LifeCamp0 (n=4 mice) shows minimal changes in delta lifetime (middle), but considerable changes in intensity (right). **e,** VTA DAN baseline response to repeated chocolate pellet consumption. Example LifeCamp delta lifetime (left, top) and intensity measurement (left, bottom) of a single animal during feeding task and averaged results across animals (middle) for LifeCamp show a change in baseline lifetime during and after feeding. Boxplots quantify comparison of LifeCamp and LifeCamp0 delta lifetime (right, top) and intensity (right, bottom) of session baseline (-4 to 0 min) with either the feeding period (0 to 10 min) or post-feeding period (10 to 15 min). LifeCamp lifetime increased during feeding (paired t-test, p=0.0148, n=6) and decreased post-feeding (paired t-test, p=0.0446, n=6). LifeCamp intensity was significantly decreased post-feeding (paired t-test, p=0.0242, n=6) but was not significantly changed during feeding (paired t-test, p=0.221, n=6) compared to baseline. LifeCamp0 lifetime was slightly decreased during feeding (paired t-test, p=00466, n=4) and unchanged post-feeding (paired t-test, p=0.0722, n=4) compared to baseline. LifeCamp0 intensity decreased during feeding (paired t-test, p=0.00465, n=4) but was not significantly different post-feeding (paired t-test, p=0.112, n=4) compared to baseline. n=number of mice. Error bars=SEM. **f,** LifeCamp intensity (left, trial baseline for normalization) and lifetime (right, session baseline for normalization) response to the first and second pellet consumption. **g,** Peak LifeCamp intensity measurements normalized to trial baseline show significant decreases in peak intensity signal of pellet 2 in comparison to pellet 1 (paired t-test, p=0.0128, n=6). LifeCamp delta lifetime measurements normalized to session baseline show no significant change in peak delta lifetime signal of pellet 2 in comparison to pellet 1 (paired t-test, p=0.0918, n=6). n=number of mice. Error bars=SEM. Boxplot elements: center mark, median; box limits, upper and lower quartiles; whiskers, furthest datapoints from the median.

Lifetime, unlike intensity, can be used to measure slow changes across long timescales; thus, we investigate whether we could utilize LifeCamp to track slow changes in intracellular [Ca] in response to feeding(33). We observed a slow increase in LifeCamp lifetime over minutes during pellet consumption (paired t-test, p=0.0148, n=6), which subsequently dropped below baseline after feeding (paired t-test, p=0.0446, n=6) (Fig. 4*E*). Although the intensity showed a similar general trend, the changes were not significant (paired t-test, p=0.221, n=6), likely due to the artifacts reflected in the LifeCamp0 intensity signal, which decreased during pellet consumption (paired t-test, p=0.00465, n=4).

Typically, intensity signals are normalized to the trial baseline by conversion to dF/F. For GECI measurements, changes in dF/F are typically interpreted as reflecting changes in action potential firing rates, ignoring potential contributions of changes in the denominator (i.e., baseline fluorescence). In contrast, lifetime can be reported and compared without normalization or by normalization to a common signal, such as the session baseline. To investigate whether baseline shifts could significantly impact dF/F intensity signals, we compared the signals during the consumption of pellet 1 with those of pellet 2. Quantification of evoked intensity changes relative to trial baseline leads to the conclusion that the magnitude of the [Ca] response to pellet 2 is smaller than that to pellet 1 (paired t-test, p=0.0128, n=6 mice) (Fig. 4*F*, *G, SI Appendix*, Fig. S5). However, lifetime measurements reported relative to session baseline show that the absolute magnitudes of the responses for pellet 1 and pellet 2 are the same (paired t-test, p=0.0918, n=6 mice). These results highlight that trial-normalized signals can be significantly impacted by baseline shifts and that lifetime, through session normalization, is insensitive to this effect.

To determine whether LifeCamp can detect differences in average Ca between neurons in different brain regions, LifeCamp was expressed in VTA DANs and in either dopamine type-1 receptor (D1R) or dopamine type-2 receptor (D2R) spiny projection neurons (SPNs) in the nucleus accumbens core (NAC) and measured in freely moving mice using FLIPR (Fig. 5*A*). We observed significantly higher baseline Ca levels in VTA DANs compared to NAC D1R- and D2R-SPNs (one way Anova, p=9.91E-12, Tukey-Kramer multiple comparison, DANs (n=5 mice) vs D1R (n=6 mice): p=2.41E-10, DANs vs D2R (n=6 mice): p=9.46E-10, n=6 mice), possibly indicating a difference in basal neuronal firing rate (Fig. 5*B*). Conversely, we observed no significant difference between D1R and D2R SPNs baseline Ca (Tukey-Kramer multiple comparison, p=0.96). The lifetime measurement of LifeCamp0 was low, consistent with no Ca activation, and did not show any difference between VTA DANs, NAC D1R-SPNs, and D2R-SPNs (one-way ANOVA, p=0.88, DANs (n=3) vs D1R (n=5) vs D2R (n=6).

**Fig. 5:**
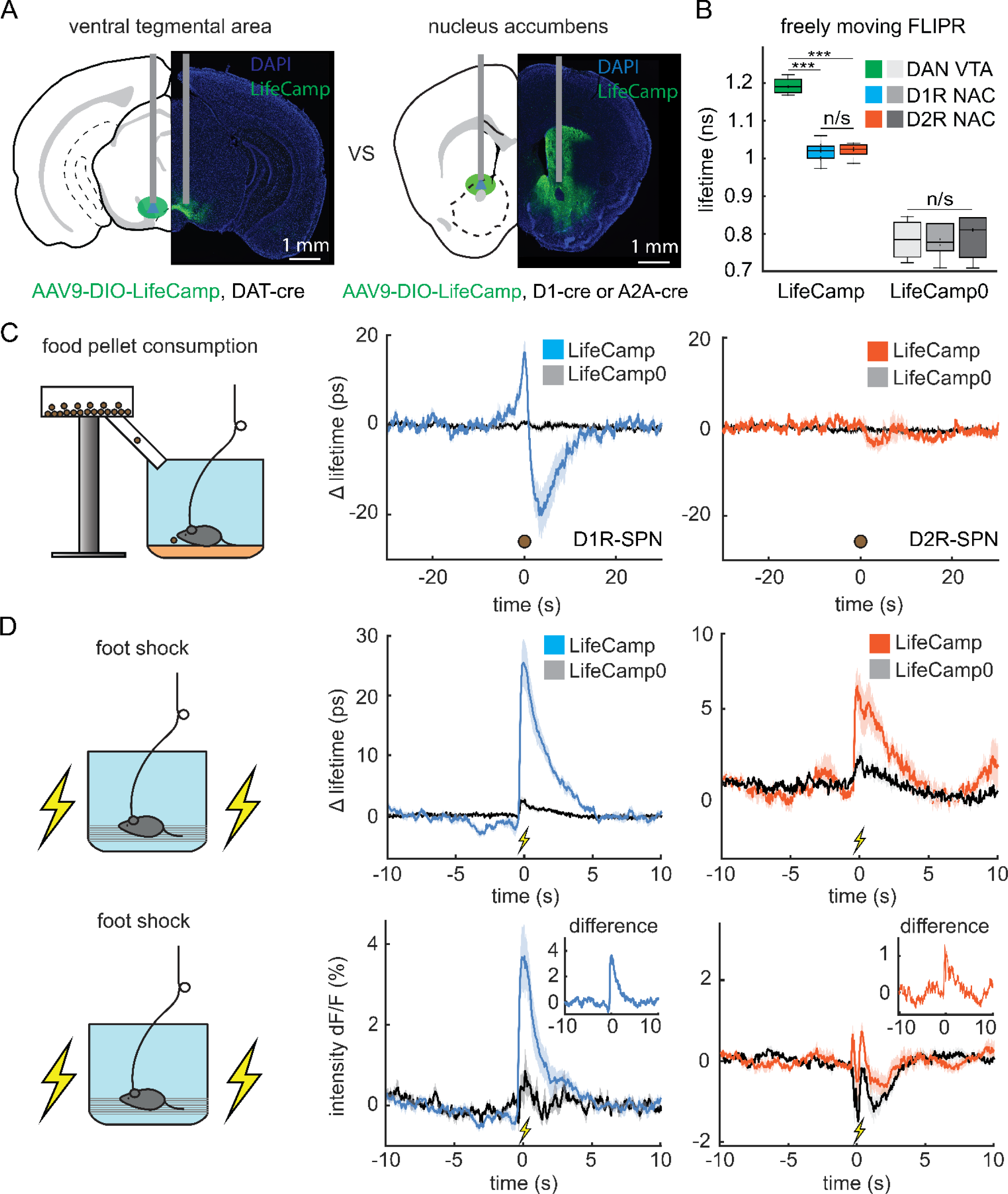
LifeCamp measured in the VTA and NAC of freely moving mice. a, Schematic and example histology of LifeCamp expression in the VTA of DAT-cre mice (left) and the NAC of D1-cre or A2A-cre mice (right), confirming expression and fiber implant location. **b,** Comparison of LifeCamp basal lifetimes between DANs and D1R- and D2R-SPNs in freely moving mice. Boxplot shows that VTA dopaminergic neurons have higher basal lifetimes than D1R-(one way Anova, p=9.91E-12, Tukey-Kramer multiple comparison, p=2.41E-10) and D2R-SPNs (Tukey-Kramer multiple comparison, p=9.46E-10) and D1R and D2R were not significantly different (Tukey-Kramer multiple comparison, p=0.96). LifeCamp0 lifetimes were not significantly different across sites (one-way Anova, p=0.88. D1R-SPN LifeCamp n=6, LifeCamp0 n=5, D2R-SPN LifeCamp n=6, LifeCamp0 n=6, DANs LifeCamp n=5, LifeCamp0 n=3, n=mice. **c,** Schematic of food pellet consumption task (left). Average delta lifetime measured response to a single trial with LifeCamp and LifeCamp0 in D1R-SPNs (middle, LifeCamp n=8, LifeCamp0 n=4) and D2R-SPNs (right, LifeCamp n=5, LifeCamp0 n=3) across animals. **d,** Schematic of foot shock task (left). LifeCamp and LifeCamp0 average delta lifetime (top) and intensity (bottom) response to foot shock in D1R-SPNs (middle) and D2R-SPNs (right). Mini plot in intensity graph (middle and right bottom) shows the average intensity LifeCamp signal subtracted by the LifeCamp0 signal. Error bar=SEM. Boxplot elements: center mark, median; box limits, upper and lower quartiles; whiskers, furthest datapoints from the median.

Activity of SPNs is difficult to measure due to their low baseline firing rate and sparsity. Thus, to determine whether LifeCamp lifetime photometry can detect changes in Ca from these challenging neurons, we expressed LifeCamp in NAC D1R- and D2R-SPNs in mice. We measured fluorescence lifetime via a fiber optic from these mice using FLIPR while giving them chocolate pellets or foot shocks. In D1R-SPNs, we observed a biphasic lifetime response to pellet consumption, whereas in D2R-SPNs, we observed minimal modulation (Fig. 5*C*). In response to foot shocks, we observed a transient increase in the lifetime signal of both D1R- and D2R-SPNs (Fig. 5*D*). LifeCamp0 lifetime was stable in both sets of mice in response to both food pellets and foot shocks.

The intensity and lifetime signals showed similar dynamics when evoked by foot shock or pellet consumption for D1R-SPNs (Fig. 5*C* and *D*, *SI Appendix*, Fig. S6*A*-*D*). However, the intensity and lifetime signals of D2R-SPNs in response to foot shocks were strikingly different (Fig. 5*D*). Intensity measurement of LifeCamp0 reported relatively large changes in dF/F during and after foot shocks and food pellet consumption, indicating the presence of artifacts, likely motion- or hemodynamic-related (Fig. 5*D* *bottom*, *SI Appendix*, Fig. S6*A*-*D*). Interestingly, subtracting the LifeCamp0 intensity signal from the LifeCamp intensity signal resulted in a trace that resembled the LifeCamp lifetime signal (Fig. 5*D*), indicating that the difference between the lifetime and intensity LifeCamp signal is likely due to artifacts in the intensity channel. Although LifeCamp0 subtraction largely restored the intensity signal, we do not recommend using this subtraction method, as artifact responses across sites may differ. Finally, we also recorded baseline Ca changes in NAC D1R-SPNs and D2R-SPNs in the foot shock task and across environments (*SI Appendix*, Fig. S7).

### Event rates and baseline calcium signals in individual neurons *in vivo*

To investigate whether LifeCamp can be used to monitor the activity of individual neurons *in vivo*, we expressed LifeCamp in cortical neurons in the primary visual cortex of mice. LifeCamp fluorescence was monitored through a cranial window and 2p-FLIM (Fig. 6*A* and *B*) in head-fixed mice on a running wheel. We observed changes in fluorescence lifetime in single neurons when firing spontaneously (Fig. 6*C*, *SI Appendix*, Fig. S8A-*C*).

**Fig. 6:**
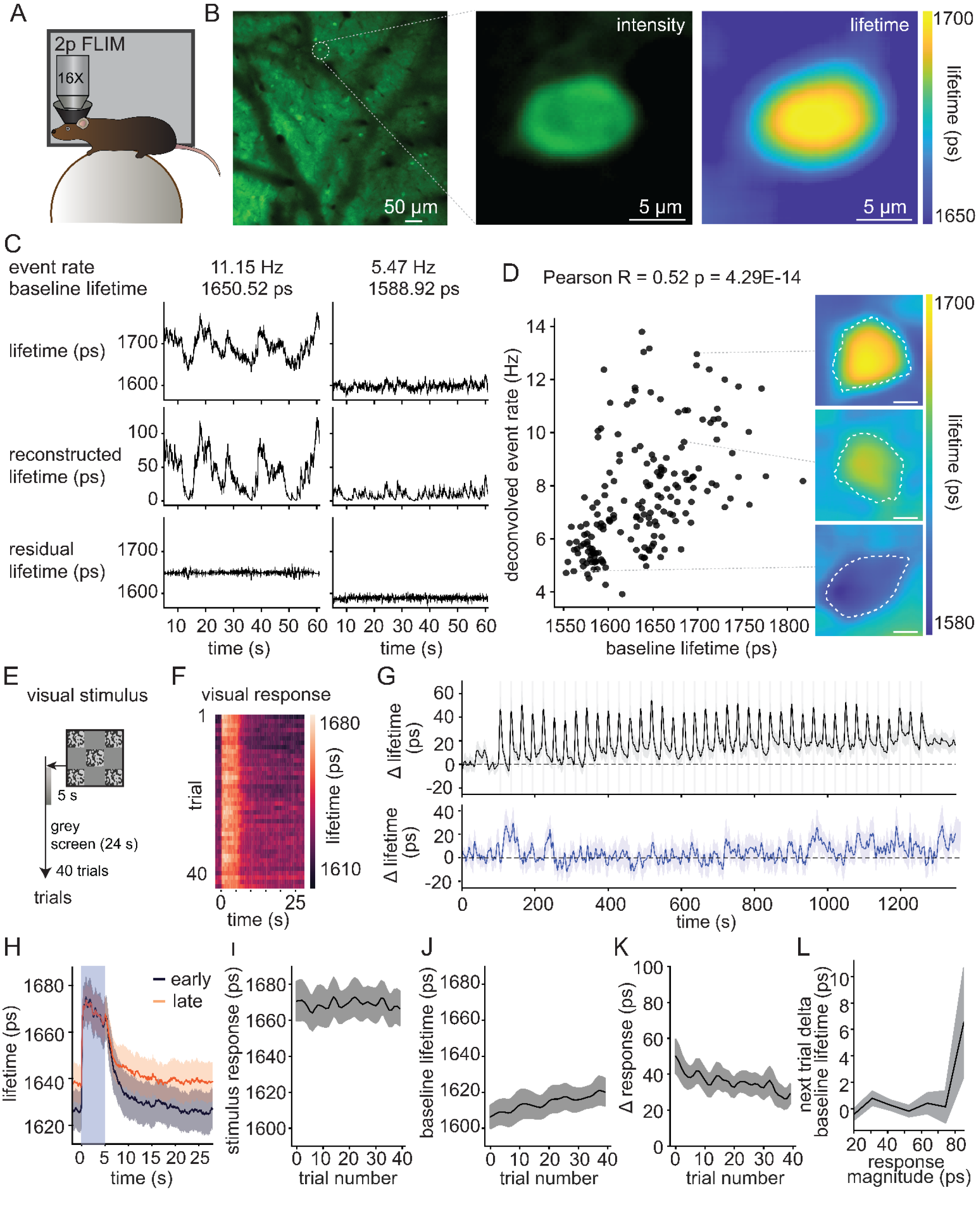
LifeCamp response signal in visual cortex neurons in mice using 2p-FLIM. a, Schematic of the *in vivo* 2p-FLIM setup used to measure from LifeCamp expressing neurons in the visual cortex. **b,** Overview (left) of the visual cortex and detailed images of a single neuron with measured intensity (middle) and fluorescence lifetime (right) using 2p-FLIM. **c,** Example neuron with higher event rate (11.15 Hz, left) and a neuron with lower event rate (5.47 Hz, right). Absolute lifetime recordings (top) were analyzed using a deconvolution algorithm, which extracted neural event rates, upon which lifetime transients were reconstructed (middle). Residual baseline lifetime (bottom) was calculated by subtracting the absolute lifetime from the event rate induced lifetime. The example neuron with larger calcium transients and higher event rate (left) has higher baseline calcium levels (lifetime=1650 ps) than the neuron with lower event rate (right) and smaller calcium transients (lifetime=1588 ps) **d,** Lifetime measurement in visual cortex shows correlation between baseline lifetime and event rate per individual neuron (*p=*4.29E-14, Pearson, R=0.524, N=180 cells, 5 mice), (left). 2p-FLIM images of three example visual cortex neurons with different measured lifetimes (right). Dotted line indicates ROI used for analysis. **e,** Schematic of visual stimulation task of 40 trials, where every trial contains 5 seconds of visual stimulation followed by an interval of 24 seconds. **f,** Example heatmap of measured fluorescence lifetime in response to visual stimulation in an example visual cortex neuron. **g,** Lifetime measured in individual neurons of visually stimulated mice (top) and unstimulated mice (bottom). Visually stimulated mice have calcium transients in response to each trial and a gradual increase in baseline calcium (*p=*9.0E-05, paired t-test between mean lifetime in pre vs. post stimulation epochs, t-statistic=4.78, n=23 cells, 4 mice). The baseline did not significantly change without visual stimulation. (*p=*0.792, paired t-test between mean lifetime in matched recording epochs, t-statistic=0.271, n=23 cells, 4 mice). Error bars=SEM. **h,** Comparison of measured average lifetime from trials early and late during the session (n=23 cells, 4 mice). **i,** Absolute LifeCamp response to visual stimulus remains stable during the session. (*p=0.0996*, Pearson R between trial number and absolute stimulus response, R=1.81E-4). **J**, Baseline calcium, calculated using the residual lifetime, increases with repeated visual stimulation (*p=*0.001, Pearson R between trial number and ITI baseline=0.108, N=23 cells, 4 mice) **k,** Delta lifetime in response to visual stimulation decreases across trials. (*p=*6.39E-05, Pearson R between trial number and stimulus delta response=-0.131) **l,** Lifetime transients evoked by visual stimulation are positively correlated with the post-trial baseline shift (*p=*0.0461, Pearson R=0.06666, N=897 trial in 23 cells). Error bar=SEM.

To investigate the relationship between baseline and transient Ca signaling in the visual cortex, LifeCamp was imaged at a high sampling frequency (∼20 Hz), and a deconvolution algorithm (OASIS) was used to identify transient events in fluorescence lifetime. We reconstructed the recorded lifetime trace from the estimated train of transient Ca events and used the residual between the initial signal and this reconstruction to determine the baseline Ca level of each neuron (Fig. 6*C*, *SI Appendix*, Fig. S8*A*-*C*). We found that cells with higher baseline [Ca] levels displayed strikingly higher Ca event rates (*p=*4.29E-14, Pearson, R=0.524, N=5 mice, 180 cells) (Fig. 6*D*, *SI Appendix*, Fig. S8D, *E*). As expected, we found that this relationship between baseline and transient Ca levels is a unique feature of the lifetime measurement, as it was not observed in the intensity measurement, which predicted similar event rates (*p=*0.223, Pearson, R=-0.0913, N=180 cells, 5 mice) (*SI Appendix*, Fig. S8*D*).

To determine whether LifeCamp can detect evoked baseline and transient Ca changes *in vivo*, head-fixed mice underwent a visual stimulation task in which they passively viewed 40 repetitions of 5-second bandpass-filtered noise movies, separated by 24-second inter-trial intervals (ITIs) (Fig. 6*E*). We observed large transient Ca responses during stimulus presentation, corresponding to a ∼50 ps change in LifeCamp lifetime. We also observed a slow increase in baseline Ca during the ITI that continued across trials (*p=*9.0E-05, paired T-test between mean lifetime in pre vs. post stimulation epochs, T-statistic=4.78, df=22, N=4 mice, 23 cells) (Fig. 6*F*, G *top* and *J*), that was not observed when visual stimuli were absent during the session (*p=*0.792, paired T-test between mean lifetime in matched recording epochs, T-statistic=0.271, df=10, N=23 cells, 4 mice) (Fig. 6*G* *bottom*).

While baseline Ca levels increased (*p=*0.001, Pearson R between trial number and ITI baseline=0.108, N=23 cells, 4 mice), absolute stimulus-evoked Ca levels remained constant over the course of repeated stimulus presentation (*p=0.0996*, Pearson R between trial number and absolute stimulus response, R=1.81E-4) (Fig. 6*H*-*J*). The delta magnitude of the stimulus-evoked transient response decreased over the course of the session (*p=*6.39E-05, Pearson R between trial number and stimulus delta response=-0.131) (Fig. 6*K*). In addition, we discovered a positive correlation between transient visually evoked calcium responses and subsequent changes in baseline lifetime (*p=*0.0461, Pearson R=0.06666, N=897 trials in 23 cells) (Fig. 6L). Altogether, this suggests that repeated visual stimulation raises baseline Ca in the visual cortex

## Discussion

We used our new rapid lifetime sensor development (RALISED) platform to create LifeCamp, a green high-speed calcium fluorescence lifetime sensor. RALISED allows for fast and rational fluorescence lifetime sensor development by linking existing intensity-based sensors to GFP variants with a different lifetime (Fig. 1*A* and *B*). LifeCamp was created by linking GCaMP8m (6), a high-speed and highly optimized calcium intensity sensor, to BrUSLEE (28), a GFP variant with low fluorescence lifetime. We show that LifeCamp allows for high-speed absolute intracellular calcium (Ca) measurements in cell culture, brain tissue, head-fixed, and freely moving mice. The excellent dynamic range of Ca-dependent LifeCamp lifetime changes (>1 ns) enables precise detection of Ca signals. We highlight new applications of LifeCamp for absolute Ca measurement, avoid existing and novel artifacts found in intensity photometry, and uncover previously unknown Ca dynamics.

### LifeCamp lifetime vs intensity calcium measurement

LifeCamp, unlike GCaMP8m, exhibits a large difference in lifetime between Ca-bound and unbound states, enabling Ca lifetime imaging, which has several advantages over conventional intensity imaging. Typical intensity measurements are sensitive to substrate-independent variables, such as sensor concentration, photobleaching, and hemodynamic/motion artifacts (6, 9–14). Therefore, Ca intensity measurements mainly allow for relative comparisons across short timescales (ms-s). In studies that use chronic imaging across days to monitor, for example, changes in cortical activity during learning, quantification is typically done by estimating Ca events and not by direct comparison of fluorescence intensity across sessions.

Lifetime imaging provides an inherently different approach. Lifetime measurements are generally insensitive to many artifacts and sensor concentration. Thus, LifeCamp enables absolute measurement of Ca levels across fast (ms-s) and slow (min-hr) timescales, facilitating easy comparison across neurons, brain regions, animals, and conditions (Fig. 4-6) (15, 18, 22, 23). When used with fluorescence lifetime microscopy (FLIM), LifeCamp enables the simultaneous measurement of event rate and baseline Ca concentration of individual neurons in brain slices and in behaving mice (Fig. 3 and 6). Our results also suggest that absolute Ca imaging could be used to screen for Ca-permeable unhealthy cells in brain slices, which could increase efficiency in electrophysiology experiments (Fig. 3*H*, *SI Appendix*, Fig. S3).

When used with fluorescence lifetime photometry at high temporal resolution (FLIPR), LifeCamp lifetime recording offers a more accurate measurement of fast changes in Ca compared to intensity measurements and slow average Ca detection (Fig. 4 and 5) (15, 18). In this study, we observed several large ligand-independent fast and slow LifeCamp fluorescence intensity artifacts, which minimally affected the simultaneously measured lifetime (Fig. 4 and 5). In some cases, for example, for slow (minutes) or small intensity signals, these artifacts entirely occluded or distorted the intensity signal (Fig. 5, *SI Appendix*, Fig. S6). Lifetime signals were minimally affected by these large swings in intensity and reflected changes in Ca, as confirmed by examining the Ca-binding mutant LifeCamp0 (Fig. 4*E* and 5*D*, *SI Appendix*, Fig. S6). In fluorescence studies, including lifetime-based measurements, a ligand-binding mutant should be used as a negative control to account for possible ligand-independent artifacts. This is especially true for the proper interpretation of small changes in lifetime, which have been made possible with sensitive approaches such as FLIPR.

Intensity signals aligned to mouse behavior are typically normalized to the trial baseline to account for slow optical artifacts such as photobleaching(9). However, this division by baseline fluorescence can produce changes in the normalized signal that reflect shifts in baseline but are erroneously interpreted as changes in the response amplitude. In our study, a comparison between the first and second food pellet consumption intensity signals (using the trial baseline for normalization) shows a reduction in the relative amplitude of the second response compared to the first. However, this difference was fully explained by the baseline shift that occurs after the first pellet consumption, as seen in the lifetime signal (Fig. 4*F*, *G*, *SI* *Appendix*, Fig. S5). This questions the validity of comparing the magnitude of intensity fiber photometry signals and concluding that they reflect differences in absolute signal levels. FLIPR LifeCamp recording circumvents this effect by detecting baseline changes, enabling more accurate interpretation of both delta and baseline Ca responses.

### LifeCamp compared to other methods

There are several methods that aim to measure absolute Ca levels within cells. Perhaps the most straightforward method for measuring Ca is to use fluorescent dyes that can diffuse into the cell (34). However, these Ca indicators do not allow for genetic targeting or easy long-term expression *in vivo*. Alternatively, several methods aim to normalize and reduce artifacts in signals from genetically-encoded intensity sensors, such as normalization to the isosbestic point (i.e., the excitation wavelength at which fluorescence is invariant to substrate concentration), normalization to an additional fluorophore of a different color, or using FRET sensors. However, the different excitation and emission wavelengths of the signal and normalization fluorescence are differentially affected by absorption, diffraction, autofluorescence, and hemodynamic changes, which can lead to incorrect correction and sensitivity to artifacts (6, 12, 14, 35, 36). Lifetime measurements circumvent these issues, as they only require a single excitation and emission channel(14). Other approaches correct for hemodynamic artifacts or normalize for sensor concentration, but often require costly additional equipment(12, 14, 37).

There has been an increased interest in the development and application of Ca lifetime sensors (38–41). However, the development of fluorescent sensors is technically challenging; indeed, the development of GCaMP8m has taken over 20 years and 8 versions to achieve high speed, signal-to-noise, and stability in biological systems(6, 42, 43). Indeed, LifeCamp achieves high speed, stability, and high signal-to-noise because it builds upon GCaMP8m. Although recently developed Ca lifetime sensors have shown excellent substrate-dependent lifetime change, they do not yet have the speed required to capture firing rates, and in some cases have not yet been shown to express in biological systems or mice (38–41). Nevertheless, these Ca lifetime sensors can be applicable to slower Ca measurements and occasions where Ca measurement is required in a different color, for example, when applying two-color FLIPR(10).

### RALISED development platform

Here, we use a new lifetime sensor development platform, termed RALISED, that enables rational, affordable, and fast development of lifetime sensors with a large dynamic range. Developing new GEFIs typically requires high-throughput screens of many constructs, high levels of expertise, expensive specialized equipment, and time (27). The key advantage of RALISED is that it bypasses the time-consuming step of new sensor optimization. By incorporating previously optimized and developed sensors, the RALISED platform allows for rapid development of new lifetime sensors at low cost and without the high levels of expertise required. This is highlighted by the fact that LifeCamp was designed from a single construct, using existing sequences, without further screening or iterative optimization. Although not critical for lifetime sensors, with RALISED it is possible to retain large intensity changes, which allows the sensor to be used for intensity and lifetime measurements.

Although not shown here, it is possible to generate new lifetime sensors using RALISED in different colors, as several static fluorescent proteins of different colors with distinctly low or high lifetime exist (28, 44–47). RALISED can also be exploited to shift the dynamic range of existing sensors: The effective ligand sensitivity of lifetime measurements is shifted to higher affinity relative to that of intensity measurements by the ratio of fluorescence intensity of the ligand bound and unbound states (methods). Finally, RALISED can be used to boost the lifetime response of existing lifetime sensors, as lifetime sensors typically also exhibit a substrate-dependent intensity change (e.g., dLight3.8). In this case, the static fluorophore should be selected so that the RALISED induced lifetime change is complementary to the inherent lifetime change of the sensor (Eq. 1).

### Fluorescence lifetime photometry using LifeCamp

Ca measurements in intensity fiber photometry studies have typically been used as a proxy for neuronal activity, as a Ca influx is triggered by each action potential (48, 49). However, cytosolic Ca levels are regulated by many other factors (5, 48). In our slice imaging experiments, neuronal firing evoked brief Ca transients, followed by a small post-firing increase in baseline Ca (Fig. 3*G*). *In vivo*, we observed clear correlations between spontaneous event rate and baseline Ca of neurons and raised Ca baseline after visual stimulation and neuronal firing (Fig. 6). In addition, using FLIPR, we observed that VTA DANs have a higher average baseline Ca compared to striatal projection neurons, which might reflect burst firing properties of DANs (Fig. 5*B*) (49–54). Altogether, these results support that slow baseline Ca population measurements using FLIPR report a combination of the average firing rate and Ca firing-dependent baseline levels that can be utilized as a proxy for neuronal activity.

In this study, we observed fast changes in Ca levels of DANs and SPNs in response to food pellet consumption and foot shock (Fig. 4*D*, 5*C* and *D*, SI Appendix, Fig. S6). Although the transient response of DANs, D1R- and D2R-SPNs to reward and punishment is known, it remains unclear how these stimuli affect slow changes in activity (55–57). Here, we observed that slow calcium levels in the VTA increased and later decreased in response to repeated food pellet consumption (Fig. 4*E*). In addition, we observed slow decreases in calcium in response to repeated foot shocks in D1R and D2R-SPNs (*SI Appendix*, Fig. S7*B*). Finally, we observed slow changes in D1R and D2R-SPN activity in response to the transition from an experimental neutral box to home cage (*SI Appendix,* Fig. S7*C*). These slow changes in Ca may reflect changes in motivation and movement(58). In addition, these results highlight that population Ca levels can be dynamically modulated across behavioral paradigms and cell types.

### Intracellular neuronal baseline calcium change

We observed changes in baseline Ca in cortical neurons in brain slices after electrical stimulation (Fig. 3*G*). In head-fixed mice, the baseline Ca and the spontaneous firing rate of individual neurons were correlated (Fig. 6*D*). In addition, we observed increases in baseline Ca of visual cortex neurons in response to visual stimulation, which correlated with the size of the transient response(Fig. 6*G*-*L*) (59). These results indicate a relationship between neuronal firing and baseline Ca levels both *in vitro* and *in vivo*, consistent with previous *in vitro* studies describing the effects of repeated stimulation or burst firing on Ca levels (60–62). The neuronal baseline Ca changes observed in this study could alter neuronal excitability, network dynamics, and synaptic transmission (63–65).

### Limitations and future directions

Here, we demonstrate the utility of LifeCamp for the measurement of cellular and neuronal Ca signals *in vitro* and *in vivo*. However, LifeCamp and the RALISED method also have limitations. For example, the lifetime range of RALISED sensors depends on the relative brightness of the intensity sensor and the static fluorophore; therefore, the effective affinity curve of the sensor depends on the excitation and emission wavelengths used, and the identity of the static fluorophore. This flexibility to tune the effective response curve can be beneficial; however, to allow for comparison across systems, the same excitation sources and emission channels need to be used. In addition, measurements with RALISED-based sensors are sensitive to possible differential bleaching rates of the sensor and the static fluorophore. Therefore, bleaching artifacts still need to be carefully considered. Lastly, RALISED relies on existing intensity sensors and cannot be used to generate sensors *de novo*. Therefore, RALISED complements but does not replace conventional sensor development.

We anticipate several future directions for the RALISED platform and the LifeCamp sensor. We foresee that RALISED will be used to develop new lifetime sensors in different colors, for example, for glutamate, GABA, and neuropeptides, and to improve the lifetime change in existing lifetime sensors. In addition, LifeCamp will enable biological investigation of the control and function of neuronal resting Ca *in vivo*, lively revealing additional targets of neuromodulators and plasticity induction.

## Methods

### Sensor design and virus production

LifeCamp was designed by connecting the available sequences of GCaMP8m and BrUSLEE with a 7x Gly-Ser-Gly (GSG) linker codon, optimized to minimize repetitive sequences(6, 28). Null mutant LifeCamp0 was designed by replacing part of the calmodulin binding site (M13 sequence) with a 5x GSG linker codon, as was previously shown to disrupt sensor responsiveness upon substrate binding(6).

To express GCaMP8m, BrUSLEE, LifeCamp, and LifeCamp0, recombinant adeno-associated viruses (AAVs) of serotype 9 or DJ/9 were used. Codons were optimized for translation in mice using the IDT codon optimization tool in Benchling. For cre-dependent expression of LifeCamp and LifeCamp0, a FLEX-cassette containing loxP and lox2272 was used. After codon optimization, constructs were produced by Twist Biosciences and cloned into a commercially available pAAV-hSyn containing plasmid (#172921, Twist Biosciences). To express constructs in HEK293T cells, the clonal gene was subcloned into a pCAG vector (Addgene, #11150), using PCR followed by Gibson assembly of each construct. Soma-targeted LifeCamp was generated by adding a RiboL1 sequence, as has been previously utilized (Addgene, #349877), but not characterized further here. For soma-targeted vectors, each construct was generated by Twist Biosciences in the pTwist Kan High Copy v2 vector (Twist Biosciences) and subcloned into a vector (Addgene, #209250). All construct sequences were validated with whole-plasmid sequencing (plasmidsaurus). AAVs were stored at −80°C. All newly generated plasmids are available through Addgene with complete sequences upon completion of submission processing (in progress): pAAV9-CAG-GCaMP8m, pAAV9-CAG-BrUSLEE, pAAV9-CAG-LifeCamp, pAAV9-hsyn-GCaMP8m, pAAV9-hsyn-LifeCamp, pAAV9-hsyn-LifeCamp0, pAAV9-hsyn-DIO-LifeCamp, pAAV9-hsyn-DIO-LifeCamp0, pAAV9-hsyn-RiboL1-LifeCamp.

### Animals and Surgery

All surgical procedures and experiments, except for the *in vivo* FLIM experiment, were performed at Harvard Medical School according to protocol approved by the Harvard Standing Committee on Animal Care. Animals, surgical procedures and experiments used for *in vivo* FLIM recordings were performed at the Beth Israel Deaconess Medical Center and approved by the Beth Israel Deaconess Medical Center Institutional Animal Care and Use committee. DAT-IRES-cre, Drd1a-cre, and Adora2a-cre mice used in this paper were bred in-house by crossing heterozygous parent lines. C57BL/6J (000664) mice were acquired from the Jackson Laboratory. Animals were provided *ad libitum* access to standard mouse chow and water for all experiments, except for those with food reward-motivated behaviors. These mice were food-restricted to 2-3g of chow per day, in addition to 10-20 20mg dustless

precision chocolate flavor pellets (F05301, Bio Serv), such that they remained at 85-95% of their initial body weight. Animals were kept under a 12/12hr dark/light cycle. All animals were older than postnatal day 56, and all experiments were performed during the dark cycle. Animals were group-housed before surgery and individually housed after surgery. To provide pain relief, oral carprofen was given to mice one day before and three days after surgery. Isoflurane was used to anesthetize mice before and during stereotactic surgery. During surgery, exposing the skull, leveling the brain, drilling burr holes in the skull, viral injection, and fiber implantation were performed using a stereotaxic frame (David Kopf Instruments). Virus was injected with a syringe pump (Harvard Apparatus, #883015) in volumes of 300 nl at a rate of 100 nL/min at the following titers: AAV9-hsyn-LifeCamp 3 X 10^12^ gc/ml or 1 X 10^13^ gc/ml, AAV9-hsyn-BrUSLEE 5.4 X 10^12^ gc/ml, AAV9-hsyn-GCaMP8m 3 X 10^12^ gc/ml or 1 X 10^13^ gc/ml, AAV9-hsyn-DIO-LifeCamp0 3 X 10^12^ gc/ml, AAV9-hsyn-DIO-LifeCamp0 3 X 10^12^ gc/ml. Mice were injected with different titers for the electrophysiology experiments: AAV9-hsyn-LifeCamp 1.5 X 10^12^ gc/ml and AAV9-hsyn-LifeCamp0 1.5 X 10^12^ gc/ml. To minimize unintentional backflow at other injection sites, pipettes rested at and above the injection site for at least 5 minutes before slowly and gradually withdrawing from the injection site.

Mice used for visual stimulation experiments underwent cranial window surgical procedures(66). 3 mm circular craniotomy was performed with a dental drill on the left lateral visual cortex. A 3 mm circular clear window (glued to a 5 mm clear window on top with edges that rest on the thinned skull) was placed onto the brain surface. The window was fixed in place with C&B Metabond (Parkell). Approximately 1 week after surgery, mice were injected 18 times at a total of 6 sites (3 depths per site: 200, 350, and 500 μm; 10-30 nl min^-1^; 33-100 nl per depth), evenly spaced throughout the exposed 3-mm-diameter circular brain surface with AAV9-hsyn-LifeCamp 1 X 10^12^ gc/ml. The window was replaced at least once a week. The following coordinates were used for stereotactic surgery (anteroposterior (AP), medio-lateral (ML) coordinates are relative to bregma, dorsoventral (DV) coordinates are relative to skull):

(VTA=ventral tegmental area, NAC: nucleus accumbens, M1: primary motor cortex, lVC: left lateral visual cortex)

VTA: -3.30 AP, +/- 0.48 ML, -4.5 DV

NAC: 1.40 AP, +/- 1.30 ML, -3.85 DV

M1: 1.0 AP, +/- 1.4 ML, -1.50 DV*

llVC: -4.6 AP, - 4.35 ML

*DV was used relative to the brain surface for injections in the M1 cortex.

All coordinates are in millimeters. Optical fibers (MFC_200/230-0.37_4.5 mm_MF1.25_FLT mono fiber optic cannula, Doric Lenses) were implanted 100-200 μm above the injection site of mice that were used for fluorescence lifetime photometry.

### Purified protein fluorescence characterization

To purify protein, His-tagged versions of BrUSLEE, LifeCamp, and GCaMP8m in bacterial expression pRsetB vectors were transformed into Escherichia coli BL21 (DE3) (New England Biolabs, C2527H), grown overnight shaking at 200 rpm at 37°C in 100 mL Luria Broth, and afterwards induced for an additional 24 hours with 0.5 mM isopropyl-β-D-thiogalactopyranoside at room temperature. The cultures were pelleted by centrifugation at 4,400 x g for 15 min, then re-suspended in lysis buffer [50 mM NaPO4, 300 mM NaCl, 10 mM imidazole, pH 8.0] supplemented with complete Mini EDTA-free protease inhibitor cocktail (Roche, 11836170001) and lysozyme. They were briefly sonicated on ice (∼50% power, 2 1-min pulses) and then supplemented with 1 μL Pierce Universal Nuclease (88701) for ∼10 min on ice. The lysate was then pelleted by centrifugation at 21,000 x g. The soluble fraction was applied to a gravity-flow Ni2+-affinity resin and washed with 10 mL of lysis buffer, followed by 10 mL of lysis buffer supplemented with 30 mM imidazole. The proteins were eluted in 5 mL of elution buffer [50 mM NaPO4, 300 mM NaCl, 500 mM imidazole, pH 8.0]. The eluate was concentrated in Amicon 10 kDa MWCO cut-off spin concentrators (Millipore, UFC5010) and then exchanged into a storage buffer [25 mM MOPS, 50 mM KCl, 50 mM NaCl, 0.5 mM MgCl2, 2.5% glycerol, pH 7.4]. Proteins were stored in the dark at 4°C until use. Protein concentrations were determined by measuring their absorbance at 280 nm prior to the assays.

LifeCamp, BrUSLEE, and GCaMP8m fluorescence lifetime was measured through time-correlated single-photon counting using a PR1 plate reader connected to an FLS980 Spectrometer. Constructs were excited using a pulsed supercontinuum laser Fiannium WhiteLaser Micro with an excitation wavelength of 480 ± 10 nm. Emission light was collected at 500 ± 20 nm, 510 ± 20 nm, and 520 ± 20 nm. All measurements were performed at 23 ± 1°C. Fluorescence lifetime was calculated using a double-exponential fitting in MATLAB 2014b, as previously described(67). LifeCamp was fitted using a custom model to determine Kd and other parameters; for details, see(10). Proteins, with a final concentration of ∼500 nM, were diluted in a buffer to 11 different concentrations of free Ca2+, ranging from 0 to 0.150 μM, and pipetted into a black, clear-bottom 96-well plate. Dilutions were made using a calcium calibration buffer kit (Invitrogen Calcium Calibration Buffer Kit #1, cat# C3008MP) with a pH of 7.2 that contained 100 mM KCl and 30 mM MOPS in deionized water. The same protein concentrations and calcium buffer solutions were used to determine the absorption and emission spectra with a fluorescence microplate reader (Synergy H1, BioTek). For the absorption spectrum measurement, the emission range was measured from 300 nm to 700 nm with 10 nm steps. For the emission spectrum measurement, the excitation range was measured from 300 nm to 700 nm with 10 nm steps.

### Two-photon fluorescence lifetime microscopy (2P-FLIM)

The intensity and lifetime of expressed proteins in HEK293T cells and cortical neurons in brain slices were measured using a custom 2-photon microscope using a Ti:Sapphire laser at 920 nm (Chameleon Vision II, 80 MHz, Coherent, Santa Clara, CA), similar to what was previously described(20). Emission light was collected using a fast photomultiplier tube (H7422-40MOD, Hamamatsu). A high-speed, 2.7 GHz vDAQ system (MBF Bioscience, Ashburn, VA) was used to collect emitted photons and the reference signal, which was controlled using ScanImage v2021.0.0 software in MATLAB 2021b to acquire single photon counting data. Data was binned at 0.391 ns for 31 bins per pulse and aligned to the reference laser pulse. Time bin 32 was used to count all photons in the entire 12.5 ns period, enabling real-time fluorescence intensity measurement. Data from HEK293T cells was collected at 1.07 Hz, imaging at 512 x 512 pixels. Lifetime data was calculated as discussed below.

### HEK293T cells culture and 2P-FLIM imaging

HEK293T cells (Lenti-X 293T Cell line, cat# 632180, Takara) were thawed and cultured in DMEM, high glucose, GlutaMAX™ Supplement, pyruvate (Gibco, cat# 10569010) supplemented with 10% Fetal bovine serum (brand) and 1% penicillin/streptomycin (Gibco, cat#20012027). To preserve the health of HEK293T cells, they were passaged before reaching 80% confluency by detaching with 0.05% trypsin-EDTA (Gibco, cat#25300054) and washing with phosphate-buffered saline (PBS) without calcium chloride and magnesium chloride (Gibco, cat#2754875) in 15 cm petri dishes. To be able to place HEK293T cells in the water bath of the 2p-FLIM set-up, cells were passaged and pipetted onto 70% EtOH precleaned coverslips in a sterile 24-well, which were either manually coated with Poly-L-Lysine (Sigma-Aldrich, cat# P4707) or ordered precoated (Corning, cat# CLS354085). 2 days later, HEK293T cells were transfected with pAAV9-CAG-GCaMP8m, pAAV9-CAG-BrUSLEE or pAAV9-CAG-LifeCamp, using transfection reagent and using the recommended protocol (TransIT-293, cat#:022431021) at room temperature. After 2 days, the coverslip containing transfected HEK293T cells was gently placed in the water bath filled with buffer solution (pH 7.5) and incubated for 20 minutes before imaging. The Buffer solution used to deplete HEK293T cells from CaCl2 contained 150 mM NaCl, 4 mM KCl, 10 mM Glucose, 10 mM HEPES, 2 mM MgCl2, 5 μM Ionomycin and 5 mM EGTA. Buffer solution to increase CaCl2 above HEK293T baseline levels contained 150 mM NaCl, 4 mM KCl, 10 mM Glucose, 10 mM HEPES, 2 mM MgCl2, 5 μM Ionomycin and either 7 or 10 mM of CaCl2. To maintain HEK293T cells at baseline levels, a buffer solution was used that contained 150 mM NaCl, 4 mM KCl, 10 mM Glucose, 10 mM HEPES, 2 mM MgCl2, 5 μM Ionomycin and 2 mM of CaCl2.

### 2p-FLIM imaging in electrically stimulated neurons

Perfusions of mice, coronal slicing, and whole-cell patch clamp recordings of primary motor cortex neurons were performed as described previously(68). Carbogen-bubbled ice-cold artificial cerebrospinal fluid (ACSF) was used to perfuse the mouse, after which it was decapitated, and the brain was extracted. Coronal slices of 300 μm were sliced using a LeicaVT1200s vibratome, after which they were incubated in a choline-based solution containing a holding chamber at 34 °C. Slices were transferred after 10 minutes into ACSF for 1 hour. ACSF contained (in mM): 125 NaCl, 2.5 KCl, 25 NaHCO3, 2 CaCl2, 1 MgCl2, 1.25 NaH2PO4, and 17 glucose (300 mOsm kg−1). Choline-based solution contained (in mM): 110 choline chloride, 25 NaHCO3, 2.5 KCl, 7 MgCl2, 0.5 CaCl2, 1.25 NaH2PO4, 25 glucose, 11.6 ascorbic acid, and 3.1 pyruvic acid. Both solutions were continuously carbon-saturated with carbogen gas (95% O2, 5% CO2). Slices were cooled down to room temperature until use. To image slices from LifeCamp and GCaMP8m expressing mice using 2p-FLIM, slices were placed in a water bath filled with ACSF held at 32-34 °C. Neurons were selected by excluding cells with a fluorophore-filled nucleus. A stimulating electrode was placed in the periphery of the selected neuron, and 2p-FLIM images were acquired at 43 Hz, at 128 x 128 pixels image size, while stimulating at a power ranging between 0.2 and 3 µA. The stimulating electrode was controlled through a custom script based on ScanImage in MATLAB 2018b. A copy of the electrode trigger pulse and a frame reference pulse were sent to a LabJack T7 data acquisition board (LabJack) and collected using LJstreamM software (LabJack) for synchronization. The Z-plane was adjusted per individual neuron to reach the brightest image plane (highest photon count) before starting a measurement and remained unchanged during recordings.

### FLIPR recordings during behavior tasks

LifeCamp and LifeCamp0 lifetime measurements in freely moving mice were performed using fluorescence lifetime photometry at high temporal resolution (FLIPR) system, as previously described(10). FLIPR lifetime and intensity signals were captured using a NI-DAQ system (NI 9215 and NI 9239, National Instruments) at 10 Khz and moving mean filtered to 10 Hz for real-time lifetime visualization. FLIPR was calibrated using coumarin 6 (2.5×10^-5^ mg/ml, Sigma Aldrich) in ethanol, following the calibration procedure as previously described(10). All behavior was captured by a camera (FL3-U3-13E4M, PointGrey). Mice were food-restricted, reaching 85-95% of their initial bodyweight. Before the start of each session, the mouse was placed in an 8 x 8-inch empty box for 10 minutes to habituate. At the start of each session, the baseline lifetime was recorded for 5 minutes, after which a 20 mg dustless precision chocolate flavor pellet (F05301, Bio Serv) was dispensed by a pellet dispenser (ENV-203-20, Med Associates Inc) every minute for 10 minutes. The pellet dispenser was controlled by a custom script in Bonsai 2.5.1 software that was synchronized with the FLIPR MATLAB 2021 software. After the last pellet was dispensed, mice were recorded for an additional 5-10 minutes. During the session, lifetime and intensity were recorded using the FLIPR system. For the foot shock experiments, mice were placed on a metal-barred floor inside a white Plexiglas box (ENV-005A, Med Associates) that delivered foot shocks controlled through a microcontroller (Arduino Uno, Arduino) programmed to run a custom script. In total, mice received 10 foot shocks (0.5 mA, 500 ms) per session, which were recorded using Bonsai 2.5.1 software. Baseline was recorded at the start of each session for 5 minutes and after the last shock was administered for 10-20 minutes. For the environment transfer experiment, mice were recorded in a 8 x 8 inch empty box (novel environment) where they were recorded for 5 minutes, after which they were gently manually transferred during the recording to their home cage and recorded for 5 minutes, after which they were placed back in the 8 x 8 inch empty box and recorded for another 5 minutes.

### 2P-FLIM visual stimulation

The experiments were performed in head-fixed mice that were able to run on a circular treadmill. Two-photon fluorescence lifetime imaging was conducted using an Ultima 2Pplus microscope controlled by Prairie View software (Bruker). Imaging was performed using a 16×/0.8 NA water-immersion objective (CFI75 LWD, Nikon) in mice implanted with a cranial window. Data were acquired at 20 frames per second with a resolution of 128 × 128 pixels per frame. A light-shielding material was used to protect the cranial window and objective from external light. XY scanning was performed using galvanometer mirrors. An InSight X3 laser (Spectra-Physics) was used to excite the fluorophores at 920 nm, and the emitted fluorescence was filtered (green channel: 525/70m-2p). The emission light was then detected by a GaAsP PMT (Hamamatsu, H11706-40) and amplified (Becker & Hickl, HFAC-26). Laser pulses and detected single-photon arrival times were time-stamped with respect to image galvanometer position (Swabian Instruments, Time Tagger Ultra). Prairie View FLIM mode was used to control all parameters, record the data, and reconstruct the Time Correlated Single Photon Counted (TCSPC) histogram image data into 256-time bins of 48 ps width. To mitigate photon pile-up, laser power was adjusted (< 15 mW) to maintain a pixelwise rate below 2–6 × 10⁶ photons per second, and the ’discard-all’ setting in Prairie View was enabled for multiple-photon-per-pulse conditions.

### Immunohistochemistry

After anesthesia with isoflurane inhalation, mice were perfused transcranially with PBS, followed by 4% paraformaldehyde (PFA) in PBS. Extracted brains were fixated in 4% PFA in PBS for 24 hours, after which they were sliced using a vibrating blade microtome (Leica Biosystems VT1000S). Slices were incubated for 2 hours in a blocking solution containing 5% normal goat serum and 0.1% Triton X-100 in PBS solution at room temperature. Then, slices were incubated overnight in primary antibody solution (1:1000, 13970, Abcam), washed in PBST (PBS supplemented with 0.1% TritonX-100), and incubated in secondary antibody containing blocking buffer (1:500, A-11039, ThermoFisher). After washing with PBST, slices were mounted onto glass slides using ProLong Diamond Antifade Mountant with DAPI (Thermo Fisher Scientific), and images were acquired using an Olympus VS200 slide scanning microscope.

### FLIM and FLIPR data analysis

2p-FLIM lifetime measurements in HEK293T cells were analyzed as follows. Lifetime was determined based on photons that were collected from 10 frames and pooled in a single image to an effective sampling rate of ∼0.1 Hz. The top 66% pixels with the highest photon counts were used for further analysis, similar to prior studies(10, 20, 69). For lifetime calculation, the selected pixels in 31 channels (or time-bins) were pooled across the image, creating a single-photon histogram with 31 bins (0.391 ns per bin). The histogram was phase-adjusted, depending on the detection of the laser sync pulse and the peak photon count of the histogram. The average lifetime was calculated using the following equation:

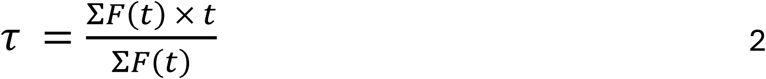

Here, F(t) is the number of photons in a given time bin (0.391 ns), with the corresponding time bin t. Lifetime was calculated per image and averaged across images to determine the lifetime value per field of view(15).

Lifetime analysis for 2p-FLIM measurements in cortical neurons was conducted similarly to the method described above, except that pixels were selected by choosing an ROI in the intensity image to isolate the individual stimulated neuron and exclude other neurons. Lifetime data was aligned to electrical stimulation by comparing reference stimulation and frame sync pulse data collected using the LabJack T7. Delta lifetime was calculated by subtracting the mean lifetime data collected from baseline (one second before stimulation) from the absolute lifetime of the trace. Responding and non-responding cells were manually characterized, and the mean of the baseline lifetime data (-1 s to 20 Hz stimulation) was used for comparison.

FLIPR analysis was performed as shown previously, see Lodder *et. al* for details(10). The FLIPR analog processing board extracted high-speed lifetime data through frequency mixing of the phase-shifted (0, 0.5, 1, and 1.5π relative to calibrated 0ns) reference pulse signals (local oscillator voltage, VLO) with the low-pass filtered (70 MHz corner frequency) photon multiplier (PMT, PMT2101, Thorlabs) signal (radio frequency voltage, VRF) to produce intermediate frequency voltages (VIF). Data was collected and processed using custom ScanImage-based data acquisition software in MATLAB 2021b. The fluorescence lifetime was calculated using Equation 3.

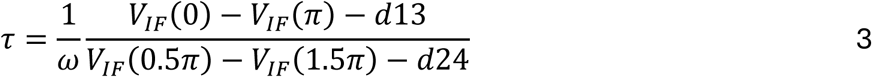

Wherein τ is the mean lifetime, ω is the angular frequency of the laser, and d13/d24 are VIF power-independent channel offsets determined during calibration(10).

Fluorescence intensity (F) is determined by collecting the direct current voltage (VDC) signal, produced by the PMT signal and filtered by the bias tee (ZFBT-4R2G+, Mini-Circuits), and by subtracting it from the VDC voltage offset (OB) determined during calibration (Eq. 4).

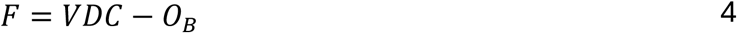

Lifetime and intensity data were averaged and down-sampled to 10 Hz or 20 Hz using a moving mean filter. Session baseline data (Fig. 4*E* *middle*, *SI Appendix*, Fig. S6*A* and *C*) was calculated using a 1-minute moving median filter. Behavioral timestamps in the food pellet consumption and the environment transfer experiment were determined using video analysis. Lifetime and intensity data were aligned using behavioral timestamps and the FLIPR synchronization signal logged by Bonsai 2.5.1 software. Baseline lifetime calculations for brain region comparison were performed by fitting an iterative Gaussian function to binned lifetime data and by extracting the mean of the fit. The first and second pellet lifetime and intensity signals that were used for comparing trial and session baseline normalization were aligned to the maximum intensity peak within 4 seconds of the behaviorally scored timestamp. Delta fluorescence lifetime average responses were calculated by subtracting the baseline (-20 to 0s for food pellet trial data, -10 to 0s for foot shock data, -4 to 0 min for session data) from the absolute lifetime signal and by calculating the mean across trials and sites. DF/F Intensity signals were calculated using equation 5.

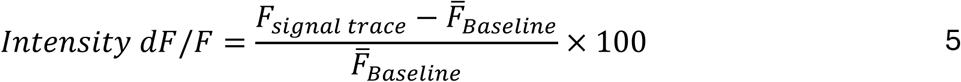

*In vivo* 2p-FLIM data were pre-processed as described previously (https://github.com/xzhang03/SPC_analysis)(70, 71). For FLIM data registration, the corresponding photon images were aligned in MATLAB using a two-step process: (1) a rigid xy-translation step, repeated three times; and (2) a non-rigid registration step using the Demonsreg function. The resulting transformations were then applied to the FLIM data. Cellpose (v3.0) was used to segment soma ROIs from the photon images(72). The fluorescence lifetime was estimated using the first-moment method, defined as the difference between the first moments of the decay histogram (mean photon arrival time) and the instrument response function (reflecting the collective response time of the imaging system). For each frame, pixels within the segmented soma ROIs were binned, and the lifetime for each ROI was estimated from the resulting binned decay histogram. For the lifetime image in Figure 6*A* and Figure 6*D*, the lifetime estimate of each pixel was calculated after local, spatial binning (bin size: 10×10 pixels, ∼4.67 um2), and the mean image of photon counts and an mean image of lifetime were plotted from the session at the resolution of the original frame (128 x 128 pixels for *in vivo* imaging). LifeCamp was imaged at a high sampling frequency (∼20 Hz), and a deconvolution algorithm (OASIS) was used to identify transient events in fluorescence lifetime.

### Modeling of lifetime data

As is well-understood, for a sensor (S) that exists in an unbound (U) and ligand (L)-bound (B) states, the fraction of sensor in the bound state is given by:

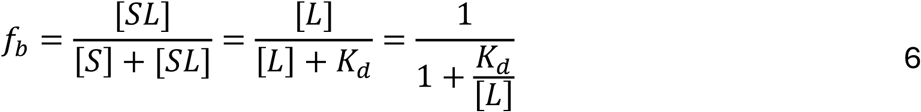

Then, under idealized conditions of no background fluorescence, the fluorescence as a function of f_b_ is given by

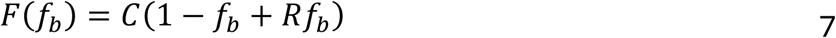

Where *R* is the ratio of the intensity of the bound vs unbound states:

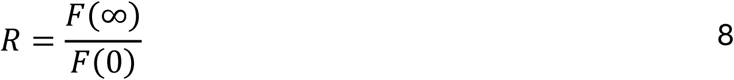

With C being a constant that takes into account many experiment-specific parameters. Then, the relationship between f_b_ and fluorescence is expressed in terms of dF/F_0_, with the denominator corresponding to the fluorescence in the unbound state *F*(*f_b_*=0):

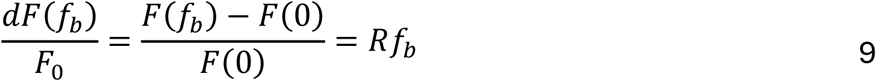

Analogously, the population lifetime tau as a function of *f_b_* is:

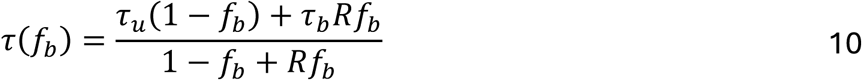

Analogous to the dF/F metric, we can define a d*τ*/*τ* metric which, in this case, is normalized to run between 0 and 1:

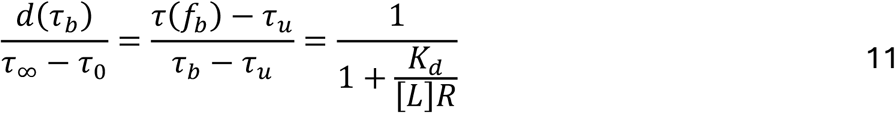

Which effectively shows that the normalized change in lifetime as a function of ligand concentration follows a typical sigmoidal activation, but with the apparent affinity shifted by a factor of R. Thus, if the bound state is brighter than the unbound state, as is the case for GCAMP and LifeCamp, then lifetime measurements are more suitable for detection of small changes.

### Quantification and statistical analysis

Data was analyzed using custom scripts in MATLAB (2021a, 2024b, and 2025a) and in Excel (v2410). Statistical sample size calculation was not performed, and experimenters were not blinded to animal groups. Paired T-test, two-sample T-test, Pearson correlation, one-way repeated measures ANOVA, and Tukey-Kramer multiple comparison statistical tests were applied to the data, as shown in the figure legends. In figures, single star (*), double star (**), and triple star (***) represent p<0.05, p<0.01, and p<0.001, respectively.

## Acknowledgements

We thank all Sabatini lab members for discussions, we thank Professor Gary Yellen for allowing us to use his lifetime plate reader, and we thank Ofer Mazor from the Harvard Medical School instrumentation core for providing patch cord holders for FLIPR calibration. We thank the Harvard Medical School Machine shop for building components of the behavioral system. This work is supported by grants to B.L.S. (NIH R35NS137336) and B.L. (PhD fellowship, Boehringer Ingelheim Funds).

