## Supplementary information for "Rapid fluorescence lifetime sensor development of LifeCamp enables transient and baseline absolute calcium measurements"

### Supplemental figures

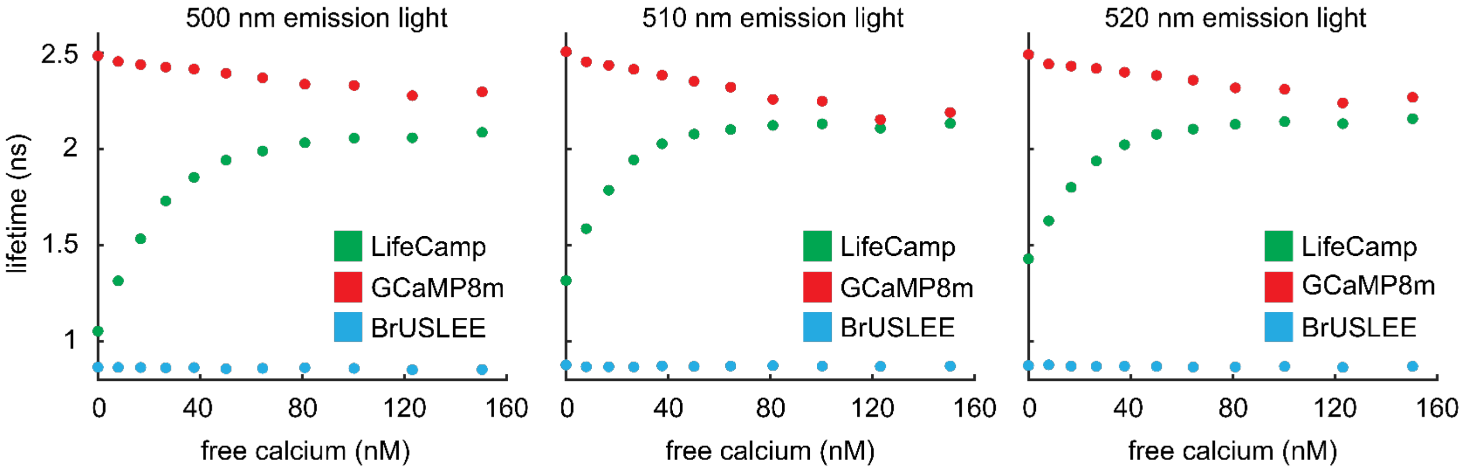


**Fig. S1: Fluorescence lifetime measured at different emission light**

Fluorescence lifetime measurement of LifeCamp, GCaMP8m and BrUSLEE using center emission wavelength of 500, 510 and 520 nm for left, middle and right graph, respectively. Constructs were exposed to different free calcium concentrations at pH 7.2 at room temperature. Lifetime was measured with 500, 510 and 520 nm (Δλ=20 nm) emission light and 480 nm (Δλ=10 nm) excitation light.

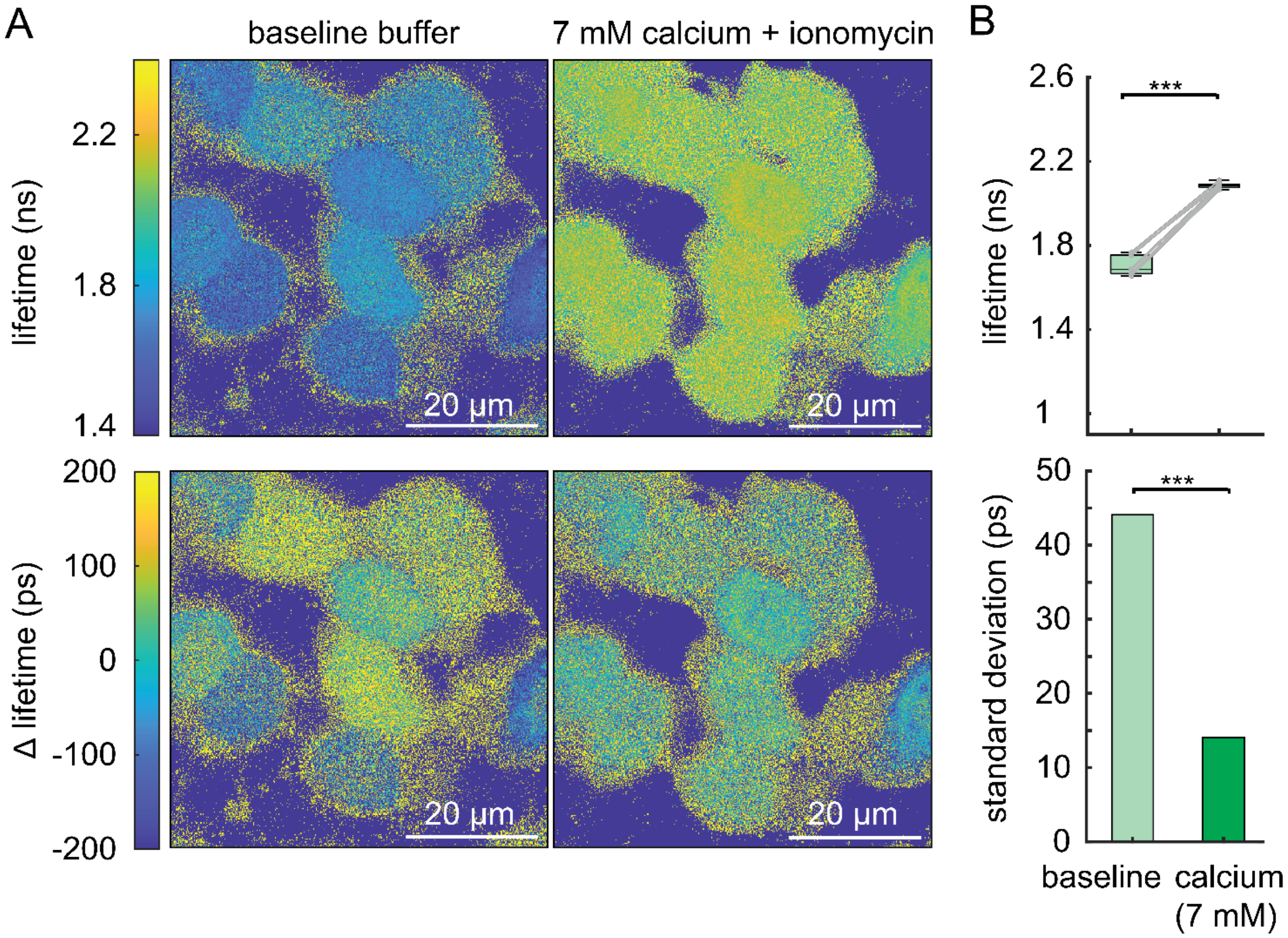


**Fig. S2: Heterogeneous intracellular calcium levels in HEK293T cells**

**a,** HEK293T cells, transfected with LifeCamp, were imaged in external bath solution with baseline [Ca] (2 mM) without calcium chelating agent (baseline condition) and in external bath solution with [Ca] (7 mM) with calcium chelating agent ionomycin (5 μM) (high calcium condition). HEK293T cells were incubated for 15 minutes prior to imaging. Top graphs show image in absolute lifetime, while bottom graphs show delta lifetime relative to image mean, to highlight differences across cells.

**b,** Fluorescence lifetime (top) of cells in baseline condition changes significantly after incubation in increased calcium condition (paired t-test, p=3.01E-13, n=12 sites). The variance (below) decreases significantly in cells that are incubated in increased calcium conditions (F-test, p=7E-4, n=12 sites). Boxplot elements: center mark, median; box limits, upper and lower quartiles; whiskers, furthest datapoints from the median.

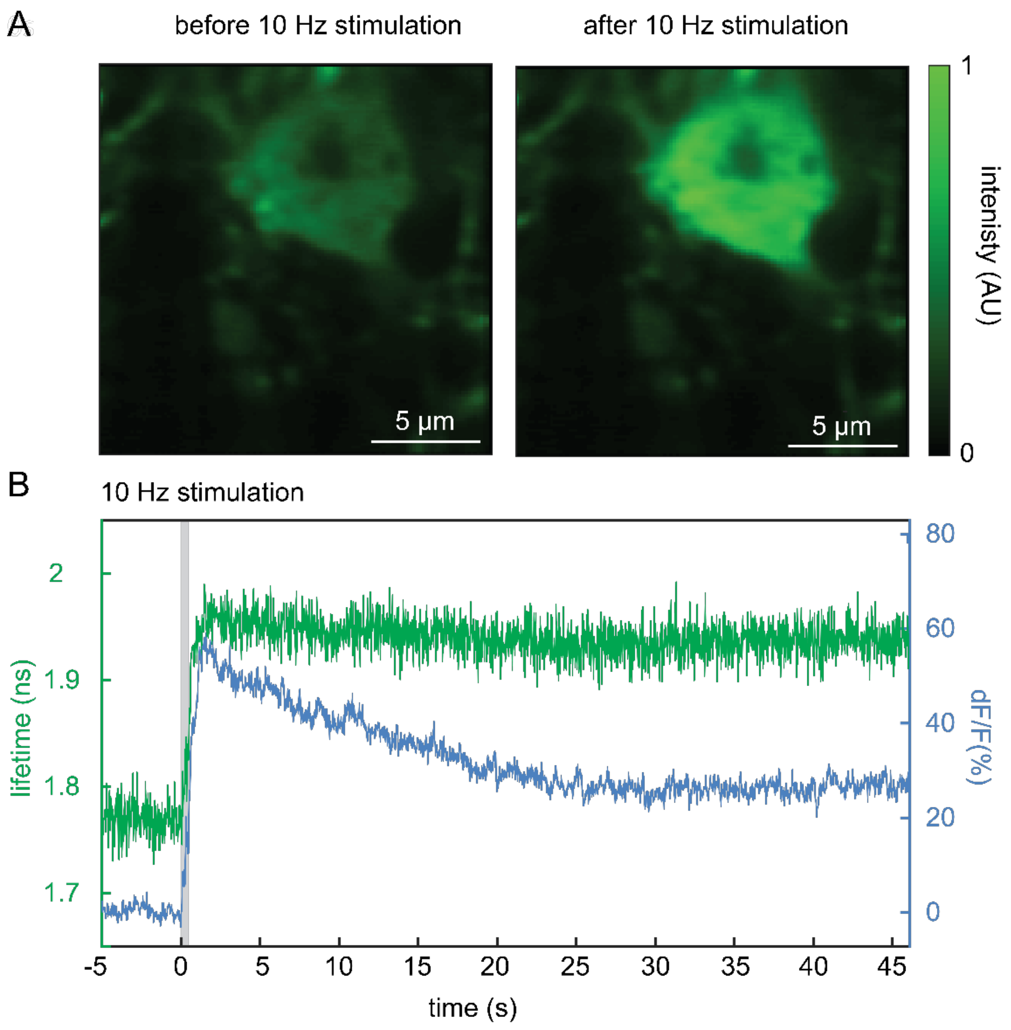


**Fig. S3: Example of individual neuron that became non-responsive after stimulation**

**a ,** 2P-FLIM image of a single neuron in the M1cortex before (left) and after it stopped responding to stimulation.

**b,** Fluorescence lifetime and intensity of an example cell expressing LifeCamp that stopped responding after stimulation at 10 Hz (grey bar).

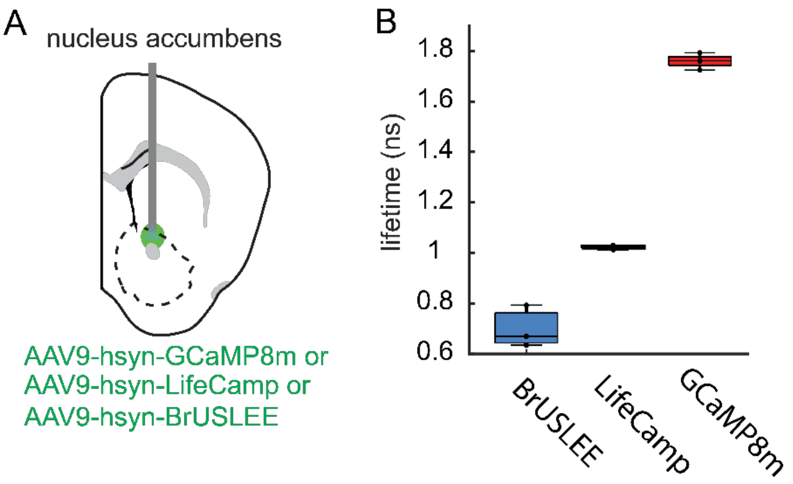


**Fig. S4: Comparison of *in vivo* lifetimes in the NAC**

**a,** Schematic of viral injection of AAV9-hsyn-GCaMP8m, LifeCamp or BrUSLEE in the NAC of mice.

**b,** *In vivo* FLIPR lifetime measurement of BrUSLEE, LifeCamp and GCaMP8m expressed in the NAC. Boxplot elements: center mark, median; box limits, upper and lower quartiles; whiskers, furthest datapoints from the median.

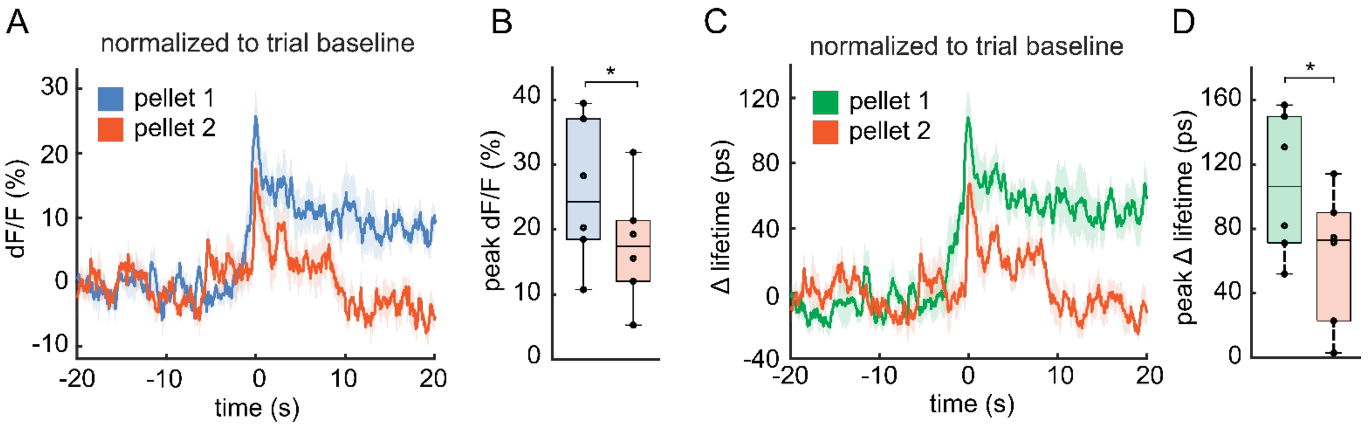


**Fig. S5: Delta lifetime signal normalization to trial baseline**

**a,** Intensity signal in response to pellet 1 and pellet 2 measured with LifeCamp relative to trial baseline.

**b** Intensity comparison of peak value in response to pellet 1 and 2 when normalized to trial baseline shows significant decreases in response signal of pellet 2 in comparison to pellet 1 ($p=0.0128$, paired t-test, $n=6$). n=number of mice. Error bar=SEM.

**c,** Delta lifetime signal from pellet 1 and pellet 2 measured with LifeCamp relative to trial baseline.

**d,** Delta lifetime comparison of peak value in response to pellet 1 and 2 when normalized to trial baseline shows decreases in response signal of pellet 2 in comparison to pellet 1 ($p=0.0175$, paired t-test, $n=6$ ). n=number of mice. Error bar=SEM. Boxplot elements: center mark, median; box limits, upper and lower quartiles; whiskers, furthest datapoints from the median.

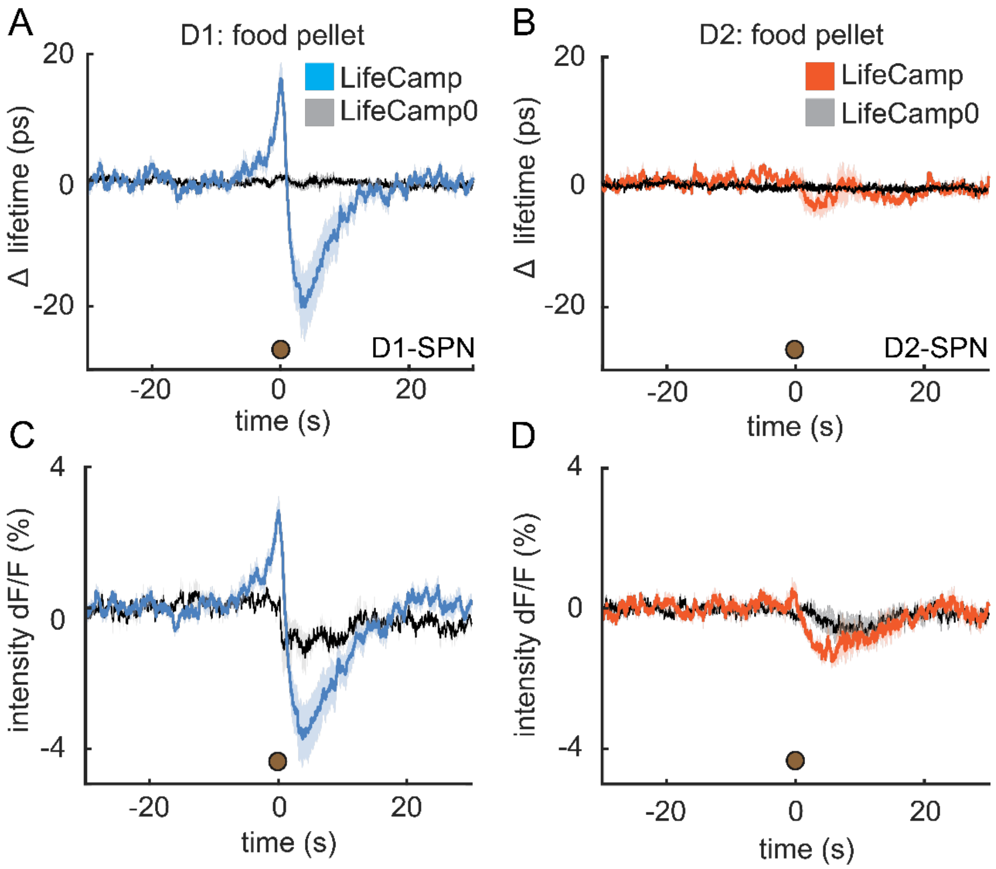


**Fig. S6: Pellet response in D1R- and D2R-SPNs**

Average LifeCamp measurements during pellet consumption show larger changes in delta lifetime and intensity in D1R-SPNs (**a, c**) compared to D2R-SPNs (**b, d**). LifeCamp0 shows changes in the Intensity signal in both D1R- and D2R-SPNs, but not in the lifetime signal.

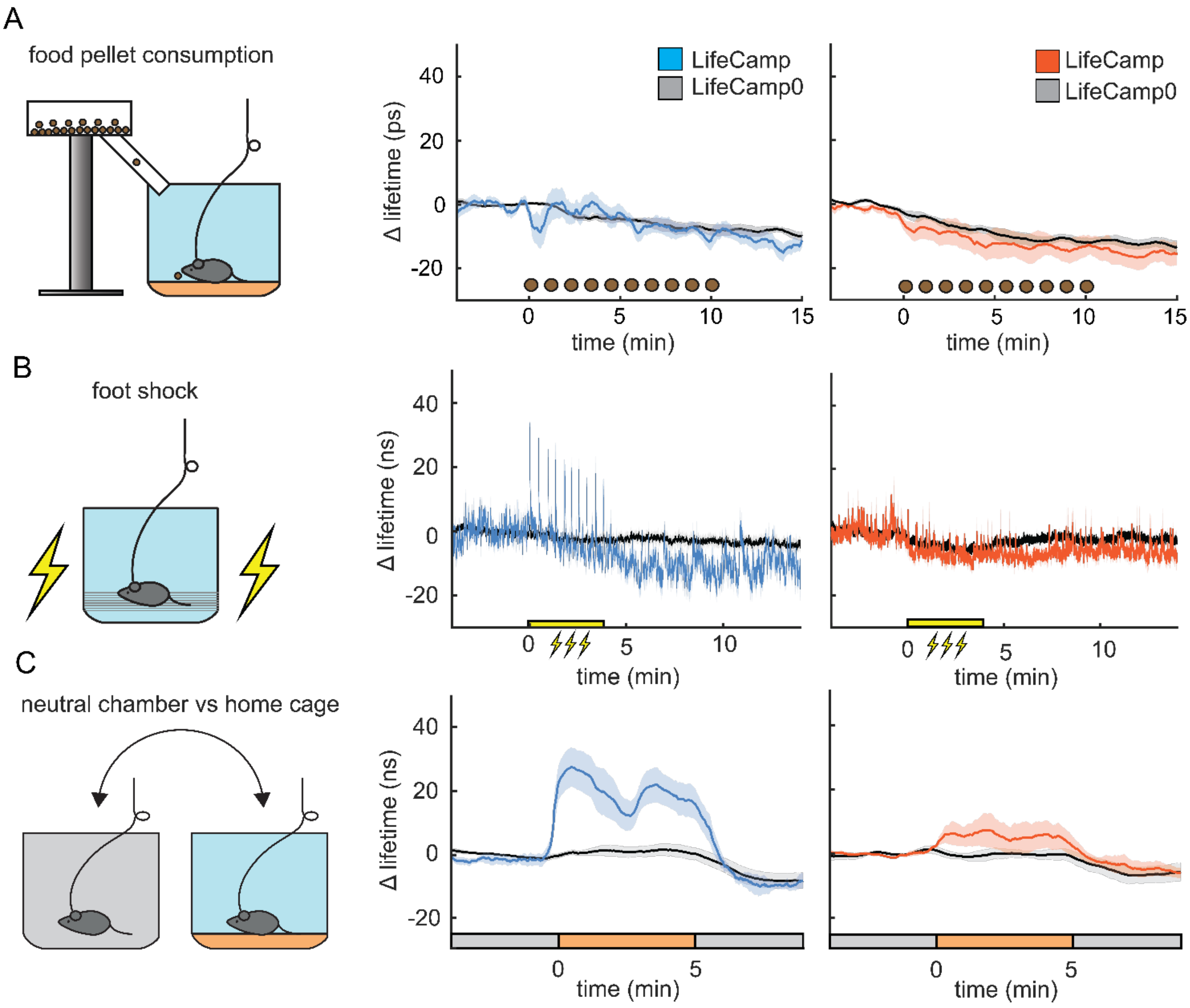


**Fig. S7: Feeding, foot shock and new environment baseline response in NAC**

**a,** Schematic of food pellet consumption task (left). Lifetime measurement during pellet consumption shows no difference between LifeCamp and LifeCamp0 during the session (middle, LifeCamp n=8, LifeCamp0 n=4) and D2R-SPNs (right, LifeCamp n=5, LifeCamp0 n=3).

**b,** Schematic of foot shock task (left). Average delta lifetime measurement of baseline in LifeCamp and LifeCamp0 during the session in D1R-SPNs (middle) and D2R-SPNs (right).

**c,** Schematic of moving mice from a neutral experimental chamber to the home cage (left). Average delta lifetime measurement shows an increased baseline in LifeCamp and not in LifeCamp0 when in the home cage in D1R-SPNs (middle) and D2R-SPNs (right) compared to the neutral experimental chamber.

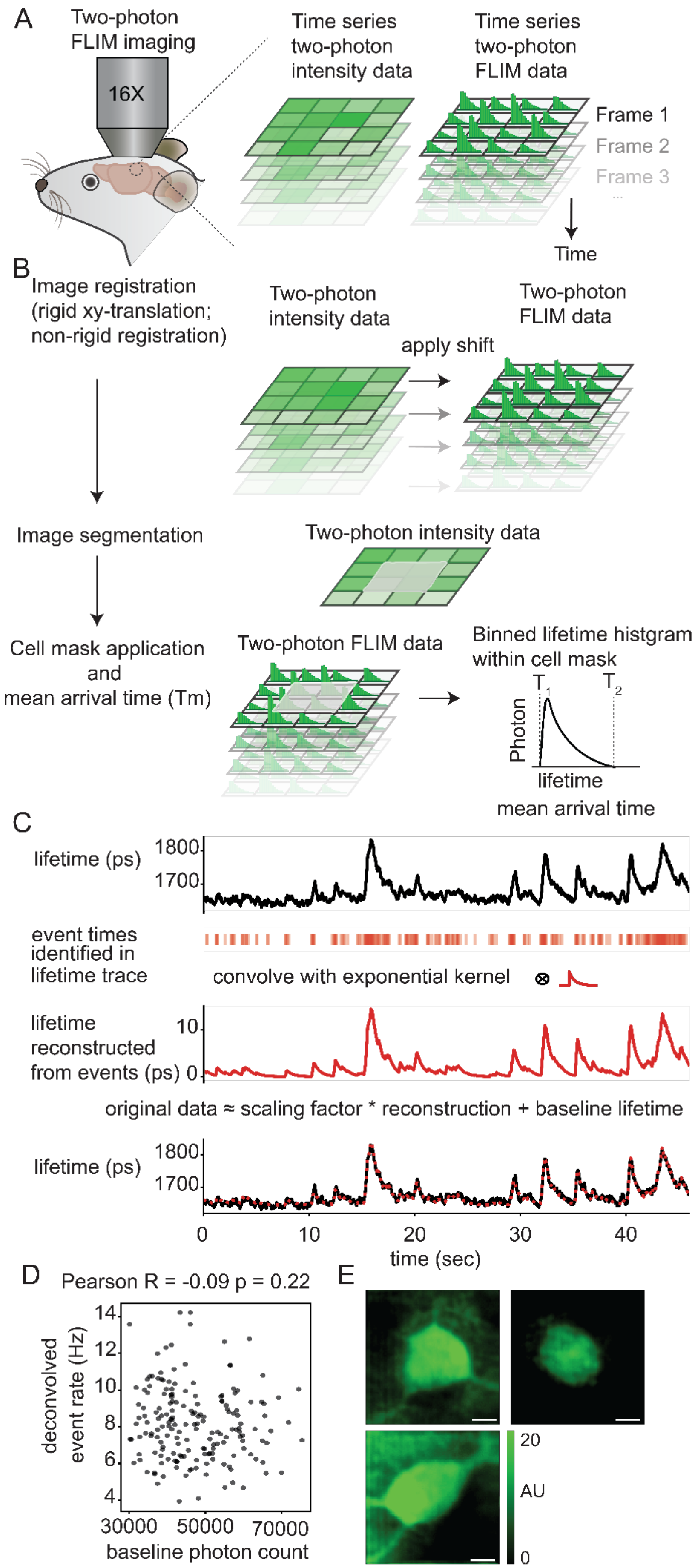


**Fig. S8: *in vivo* 2p-FLIM method and intensity recording**

**a,** Schematic of 2p-FLIM imaging setup in head-fixed mice to measure and calculate fluorescence lifetimes of LifeCamp expressing visual cortex neurons.

**b,** Schematic of fluorescence lifetime image processing.

**c,** Example lifetime trace of a visual cortex neuron, identified event times in lifetime trace, exponential kernel used to convolve absolute lifetime data, lifetime reconstructed from events, overlay of reconstructed lifetime trace on top of original data used to identify residual baseline data (top to bottom).

**d,** Baseline photon count was not correlated with deconvolved event rate. Every dot represents a single neuron. ($p$=0.223, Pearson, R=-0.0913, N=180 cells, 5 mice),

**e,** intensity images of example neurons imaged using 2p-FLIM.
